# Membrane thickness, lipid phase and sterol type are determining factors in the permeability of membranes to small solutes

**DOI:** 10.1101/2021.07.16.452599

**Authors:** Jacopo Frallicciardi, Josef Melcr, Pareskevi Siginou, Siewert J. Marrink, Bert Poolman

## Abstract

Cell membranes provide a selective semi-permeable barrier to the passive transport of molecules. This property differs greatly between organisms. While the cytoplasmic membrane of bacterial cells is highly permeable for weak acids and glycerol, yeasts can maintain large concentration gradients. Here we show that such differences can arise from the physical state of the plasma membrane. By combining stopped-flow kinetic measurements with molecular dynamics simulations, we performed a systematic analysis of the permeability through synthetic lipid membranes to obtain detailed molecular insight into the permeation mechanisms. While membrane thickness is an important parameter for the permeability through fluid membranes, the largest differences occur when the membranes transit from the liquid-disordered to liquid-ordered and/or to gel state. By comparing our results with *in vivo* measurements from yeast, we conclude that the yeast membrane exists in a highly ordered and rigid state, which is comparable to synthetic saturated DPPC-sterol membranes.

## Introduction

Membrane permeability is an essential property of cell membranes which regulates the passage of solutes and solvents into and out of cells or intracellular compartments. Permeation is either mediated by membrane transport proteins or channels or occurs via passive diffusion; here we focus on the latter. Biological membranes are relatively impermeable for most nutrients and ions, hence passive permeability contributes to the flux of a subset of molecules.

Passive diffusion is dependent on the molecular characteristics of both the solute and the lipid bilayer. While most literature focuses on the properties of the permeants, namely size, shape, and polarity^1–6^, fewer studies tackle the effects of membrane lipid composition on passive diffusion. According to the solubility-diffusion model, the permeability coefficient (*P*) of a molecule is predicted to be inversely proportional to the thickness of the membrane, and proportional to the product of partition coefficient and diffusion coefficient in the membrane. As a proxy for the partitioning coefficient of a compound in the membrane, partitioning into organic solvents is traditionally considered. Indeed, a strong correlation of permeability with the octanol-water partition coefficient has been found in egg lecithin membranes, with the permeability varying over five to six orders of magnitude^7–9^. In numerous works, however, highly ordered membranes have been shown to exhibit permeability coefficients significantly different from those predicted by the solubility-diffusion model using the octanol-water partition coefficient for solubility^10–13^. Despite many studies on the passive permeability of biological membranes, there is no data that addresses systematically the role of lipid composition on the permeability of membranes for small molecules.

In our previous work^14,15^, we developed an assay and analysis method to accurately determine the permeability coefficient of synthetic membranes and the plasma membrane of living cells for small molecules such as weak acids and bases, water, and neutral solutes. The vesicles or cells are osmotically shocked by the permeant and the rate of volume recovery or cytosolic acidification/alkalinization is analyzed. By testing vesicles of different lipid composition, we observed a relationship between acyl chain saturation and membrane permeability for both water and formic and lactic acid.

Here, we combine the experimental work with coarse-grained (CG) molecular dynamics (MD) simulations to unravel the determinants of small molecule permeability of lipid membranes. MD simulations have previously revealed that the permeability of small molecules depends not only on their chemical nature, but also on the membrane properties^5,9,16,17^. For instance, the permeability of water molecules through model liquid-ordered (L_O_) membranes is lower than through model liquid-disordered (L_d_) membranes^18^. Moreover, as is also shown in^5^, this study reveals why the solubility diffusion model breaks down: it does not capture the differences between the two membranes well when the compound solubility is assumed to be the same in the two membrane environments as in the commonly used octanol. This difference becomes increasingly important in membranes with more complicated compositions and rich phase behavior like the plasma membrane of yeast, which contains not only patches of highly ordered lipids in the L_O_ phase, but also regions in the gel (L_β_) phase^19–22^. Such details can be obtained from MD simulations, in particular using a CG model which enable systematic exploration of membrane composition and state points^23,24^, allowing us to find the main descriptors of the permeability through biomembranes.

Using the combined approach of MD simulations and experimental fluorescence measurements we determine the permeability coefficients of several polar compounds (weak acids and glycerol) through membranes of different lipid compositions. The compositions span varying saturation levels, acyl chain lengths and sterol concentrations with a special emphasis on membranes in different physical states. By comparing the experiments with the results from MD simulations, we are able to interpret the observed changes in terms of solubility and solvent proximity. Finally, we compare the resulting permeability coefficients with the values we estimated *in vivo* for *Saccharomyces cerevisiae*^14^ and use the permeability measurements as indirect reporters of the physical state of the yeast plasma membrane.

## Results

### Permeability and partitioning coefficients from experiments and simulations

In experiments, we determine the permeability coefficients using a fluorescence-based assay that reports volume changes of vesicles by means of calcein self-quenching fluorescence^14^. Briefly, we follow the out-of-equilibrium relaxation kinetics of vesicles upon osmotic upshift with a stopped-flow apparatus, by addition of an osmolyte to the vesicle solution. The thermodynamic equilibrium is re-established by the flux of water and/or the osmolyte. The contribution of the two fluxes to the recovery kinetics depends on the relative permeability of water and the osmolyte. The permeability coefficients are then calculated using the previously developed technique and analysis tool^14,15^. Measured vesicle volume relaxation curves from solutes that permeate with slow or fast kinetics (formic acid, L-lactic acid, glycerol), and from non-permeating solutes like KCl, are shown in Fig. 1A. In the case of the weak acids, the vesicle volume recovers only partially because the acids were added as sodium salts, and sodium ions do not permeate through lipid membranes on the timescale of the measurements. In contrast, glycerol leads to an overall inflation of the vesicles above their original volume after the concentrations in/out of the vesicles are equilibrated. This is caused by the preferential partitioning of glycerol to the membrane-water interface and its interactions with lipids.

**Figure 1.**
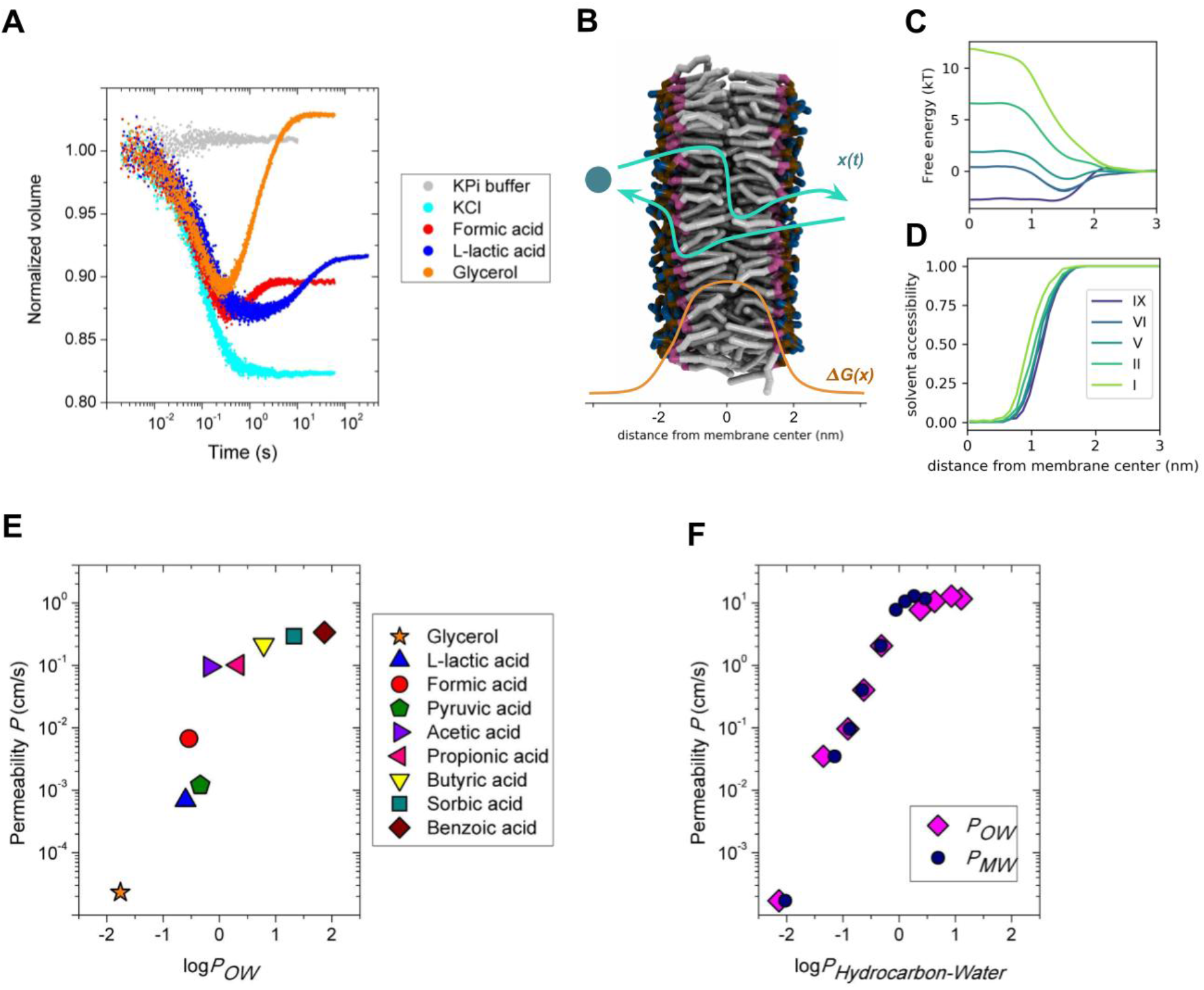
A. Overview of experimental assay. Kinetic data obtained with the calcein self-quenching assay using vesicles composed of DOPC mixed with buffer (grey) or osmotically shocked with 52.5 mM KCl (cyan), 50 mM sodium formate (red), 50 mM sodium L-lactate (blue) or 120 mM glycerol (orange). B. Schematic description of the permeation process, *x(t)*, through a lipid membrane with an example free energy profile, *ΔG(x)* (lipid tails, grey; glycerol moiety, purple; phosphate moiety, ochre; and choline moiety, blue; water molecules are not shown). C. Selected free energy profiles from simulations of solutes with varying hydrophobicity levels (I most hydrophilic, IX most hydrophobic) permeating through a DOPC lipid membrane as a function of the distance from the bilayer center along the membrane. Only one half of the whole symmetric permeation profile is shown. D. Solvent accessibility profiles of the permeating solutes along the permeation pathway. Solutes interact with solvent molecules even deep in the membrane tail region (x<1.0 nm). E, F. Permeability coefficients for DOPC membranes from experiments (E) and simulations (F) plotted against the octanol/water and membrane/water partitioning coefficient (P*_OW_* and P*_MW_*) of the solutes. The numerical values are presented in Supplementary Table 1.

In MD simulations, we have used the coarse-grained (CG) Martini 3 model^25^ to describe the permeation of a range of hydrophobic and hydrophilic generic solutes at infinite dilution, using the inhomogeneous solubility-diffusion model in which the permeation rate is obtained as an integral across the membrane of the ratio between local solubility and friction (see Methods, Eq. 1). Briefly, we obtain free energy profiles of the solutes as a function of the distance from the membrane center (linked to the local solubility via the Boltzmann factor) by sampling its translocation through the membrane as schematically described in Fig. 1B. The solutes are modeled as single CG beads and labeled according to their hydrophilicity with levels I-IX. The most hydrophobic particles, levels VIII and IX, have log*P_OW_* between butyric and sorbic acids, level V represents the hydrophobicity of pyruvic acid, and the most hydrophilic particle, level I, has log*P_OW_* comparable to phosphoric acid. The free energy profiles through a DOPC membrane of solutes with different octanol-water partition coefficients, log*P_OW_*, and their solvent accessibility are presented in Fig. 1C and D, respectively. Profiles of local friction are in Supplementary Fig. S1.

We compare both methods by plotting the dependence of the obtained permeability coefficients, *P*, on the octanol-water partitioning coefficients, log*P_OW_*, of several hydrophobic and hydrophilic solutes (eight weak acids and glycerol) in Fig. 1E and F (numerical values in Supplementary Table 1). The permeability coefficients from MD simulations are somewhat larger than in experiments because of the intrinsically faster dynamics in coarse-grained modeling, but the relative changes of the permeability coefficients with log*P_OW_* are in good accordance with experiments. For instance, the slope of the dependence of *P* on log*P_OW_* for the hydrophobic and hydrophilic solutes is independently recovered in both methods. In particular, the difference between acetic acid (log*P_OW_* = −0.17) and formic acid (log*P_OW_* = −0.54) corresponds to approximately one order of magnitude change of their permeability, whereas a similar difference from propionic acid (log*P_OW_* = 0.33) to butyric acid (log*P_OW_* = 0.79) is reflected only by about a factor of 2 increase of the permeability. Interestingly, the membrane-water partition coefficient, log*P_MW_*, from MD simulations is smaller than log*P_OW_* for the hydrophobic compounds (Fig. 1F). This is caused by non-vanishing interactions of the solutes with solvent molecules even below 1 nm from the membrane center (Fig. 1D), making the overall polarity of the lipid membrane interior generally higher than in bulk octanol.

### Phospholipids tail length affects permeability by changing the membrane thickness

We determined permeability coefficients through phospholipid bilayers of different thickness with mono-unsaturated tails of lengths between 14 and 26 carbons (Fig. 2A and 2B). The permeability coefficients for formic acid, L-lactic acid and water decrease with increasing chain length (Fig. 2A), and they are in line with the calculated values from simulations using the solute of hydrophobic level III as a representative for a generic polar weak acid in neutral form (Fig. 2B). Increasing the length of the phospholipid acyl tails increases the hydrophobic thickness of the membrane (Fig. 2B). Elongating the lipid tails by 4 carbons decreases the permeability coefficients by approximately one half for tail lengths from 14 to 26 carbons.

**Figure 2.**
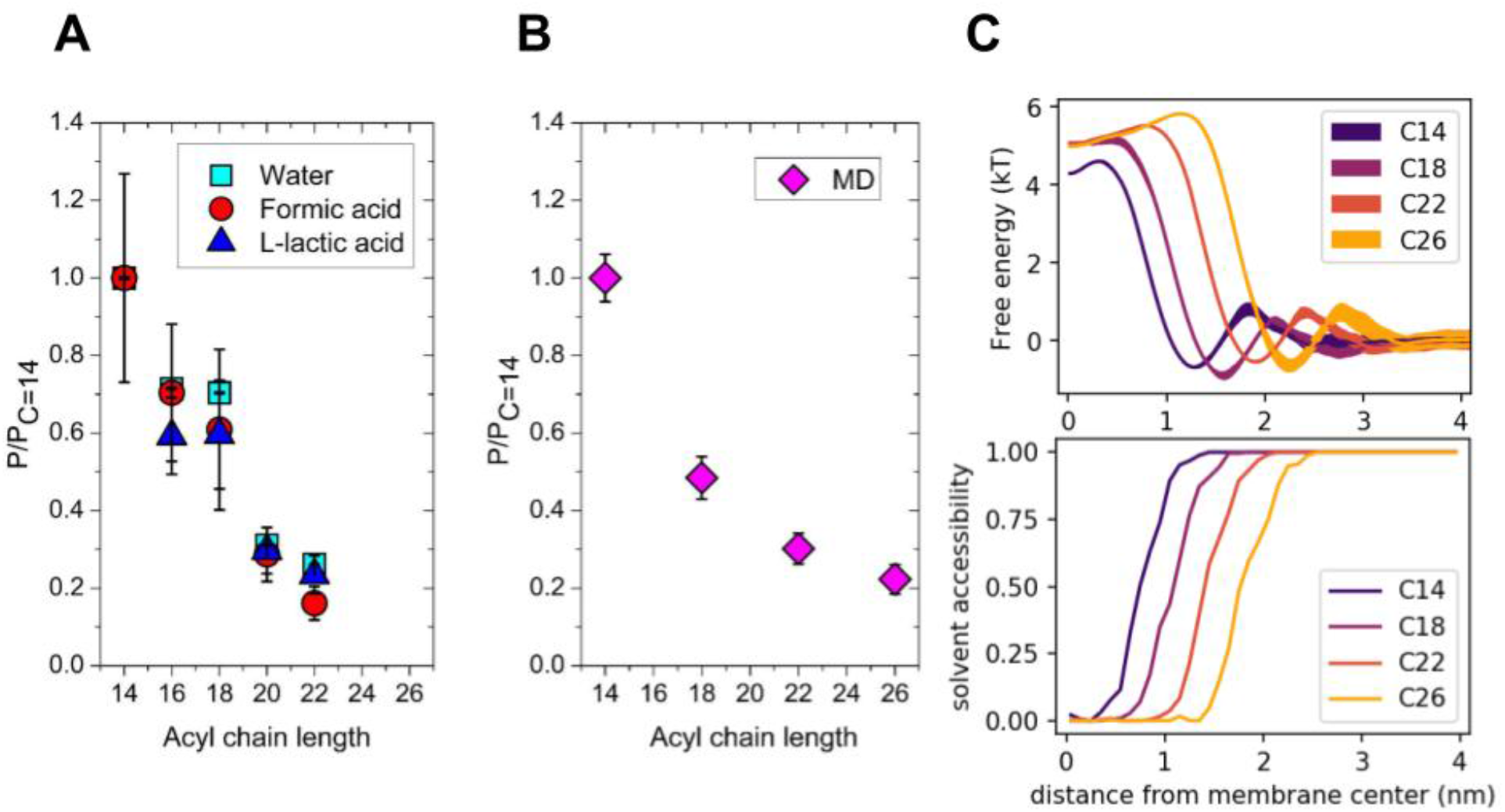
Permeability of solutes as a function of acyl tail length from experiments (A) and simulations (B). Permeability coefficients are normalized to *P_C=14_*, which in the experimental analysis corresponds to 22.8 (±1.9) x 10^-3^ cm/s, 11.80 (±1.43) x 10^-3^ cm/s, and 0.332 (±0.052) x 10^-3^ cm/s for water, formic acid, and L-lactic acid, respectively. The numerical values are presented in Supplementary Table 2. C. Free energy (top) and solvent accessibility (bottom) profiles from simulated membranes of variable thickness. The uncertainty of the free energy profiles is represented by the thickness of the lines. Longer acyl chain length of the phospholipid tails increases the membrane hydrophobic thickness, which leads to an increasingly wider and higher free energy profile.

A higher membrane thickness leads to a net shift in the position of the membrane interface with water (Fig. 2C bottom). The free energy barrier shows corresponding net shifts of the profiles, yielding wider barriers for thicker membranes (Fig. 2C top). Moreover, the free energy barriers reveal a decreasing height for thinner membranes. This is linked both to the overall increased polarity of the membrane interior for thin bilayers with respect to thick bilayers, as the solvent molecules are closer to the membrane center for very thin bilayers (*i.e*., C14 in Fig. 2C), and to the changes of the lipid packing as reflected in the changes of average lipid surface areas (Supplementary Fig. S2).

### Membrane phase transition from L_d_ to L_β_ decreases the permeability of the membrane by orders of magnitude

We have measured permeability coefficients of water, L-lactic acid, formic acid, and glycerol through lipid membranes with a degree of unsaturation, *d*, from 1 (DOPC) to 0 (DPPC); here we also measured glycerol as the plasma membranes of examined bacteria and yeast have orders of magnitude difference in permeability for this osmolyte. From both our measurements and MD simulations, we observe that decreasing the degree of unsaturation of the phospholipid acyl chains from 1 to 0.17 leads to a corresponding gradual decrease of the permeability coefficient for all the compounds (Fig. 3A and 3B). The decrease of the permeability coefficients arises from wider and higher free energy profiles as shown in Fig. 3C. In line with previous work^14^, the effects of the degree of unsaturation on the permeability coefficient are almost independent of the chemical nature of the permeants in the range of *d* between 1.0 and 0.17 (Fig 3A).

**Figure 3.**
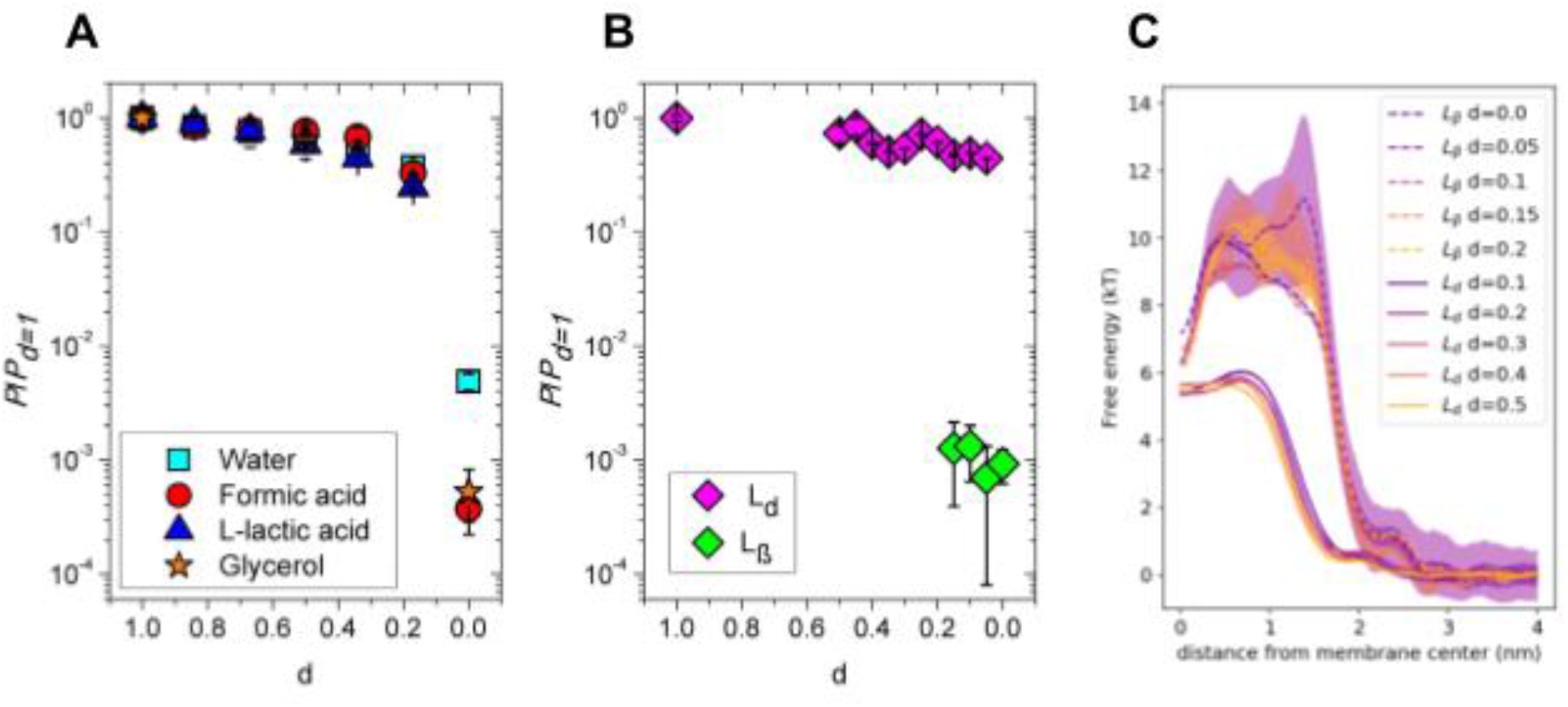
Permeability coefficients as a function of acyl tail unsaturation (*d*) for bilayers with PC headgroup. The permeability coefficients for each solute are normalized to their respective value at *d*=1.0, which in the experimental analysis corresponds to 16.0 (±1.7) x 10^-3^ cm/s, 6.73 (±0.74) x 10^-3^ cm/s, 0.198 (±0.033) x 10^-3^ cm/s, and 2.3 (±0.2) x 10^-6^ cm/s for water, formic acid, L-lactic acid, and glycerol, respectively. Permeability coefficients for water (cyan squares), formic acid (red circles), L-lactic acid (blue triangles), glycerol (orange stars) from experiments (A) and from MD simulations (B) for membranes in the L_d_ and L_β_ phase. Simulations in the range of *d* between 0.17 and 0.05 show results from both membrane phases, L_d_ and L_β_, which were used as a starting configuration and remained meta-stable within the simulation time. The numerical values are presented in Supplementary Table 3. C. Free energy profiles from simulated membranes at L_d_ phase (solid lines) and L_β_ phase (dashed) with varying degree of unsaturation *d*.

Importantly, for membranes with a degree of unsaturation below *d*=0.17, we observe very dramatic changes of the permeability coefficients. Namely, the permeability coefficient through DOPC (*d*=1.0) versus DPPC (d=0.0) membranes decreases approximately 200-fold for water, and 2000-fold for formic acid and glycerol. Moreover, permeation of lactic acid was not observed in the DPPC vesicles on the timescale of 10 hours.

The enormous leap in the permeability coefficient arises mainly from the highly decreased solubility of the compounds in the membranes with low degrees of unsaturation as seen in Fig. 3C. The free energy profiles from MD simulations show not only a wider but also much higher barriers, which form the major contribution to the decrease of the permeability coefficients. In addition, the mobility of the permeating solutes is significantly decreased slowing the permeation even further, however, this effect is lower for smaller solutes (Supplementary Fig. 3).

The sudden non-smooth changes of membrane properties, including the differences in permeability coefficient, are directly linked to the phase state of the membranes. In Fig. 3B, we plot the calculated permeability coefficients from simulations for different phases, that are, L_d_ and L_β_. The membrane phase state remains *meta-stable* on the simulation time scales, giving rise to two distinct values for the permeability coefficient in the region of *d* between 0.15 and 0.05. While the free energy and friction profiles from the membrane at the L_d_ phase compare well to those of the POPC membrane (L_d_ phase under the same conditions), the profiles from membranes at the L_β_ phase are similar to those of DPPC membranes (L_β_ phase).

We corroborate our findings by DSC measurements (Supplementary Fig. 4), which show a decreasing membrane melting temperature with increasing degree of unsaturation. In particular, the POPC/DPPC mixtures with d=0.17 and d=0.34 form a stable L_d_ phase with an interface to the L_β_ phase through possible coexistence (Supplementary Fig. 4A); note the broad transition peaks around 24°C (d = 0.34) and 34°C (d = 0.17). Phase coexistence has been previously reported in GUVs prepared from the same lipid species by fluorescence measurements using probe partitioning^26^.

### Sterols modulate the solute permeability by affecting the membrane phase state

We have selected cholesterol and ergosterol as the main sterols of mammalian and yeast plasma membranes^27,28^, and assessed their effects on the membrane permeability at molar levels from 0 to 45%. Permeability coefficients of membranes with varying concentrations of sterols and unsaturation index from experiments and simulations are shown in Fig. 4, and all numerical values are given in Supplementary Table 4.

**Figure 4.**
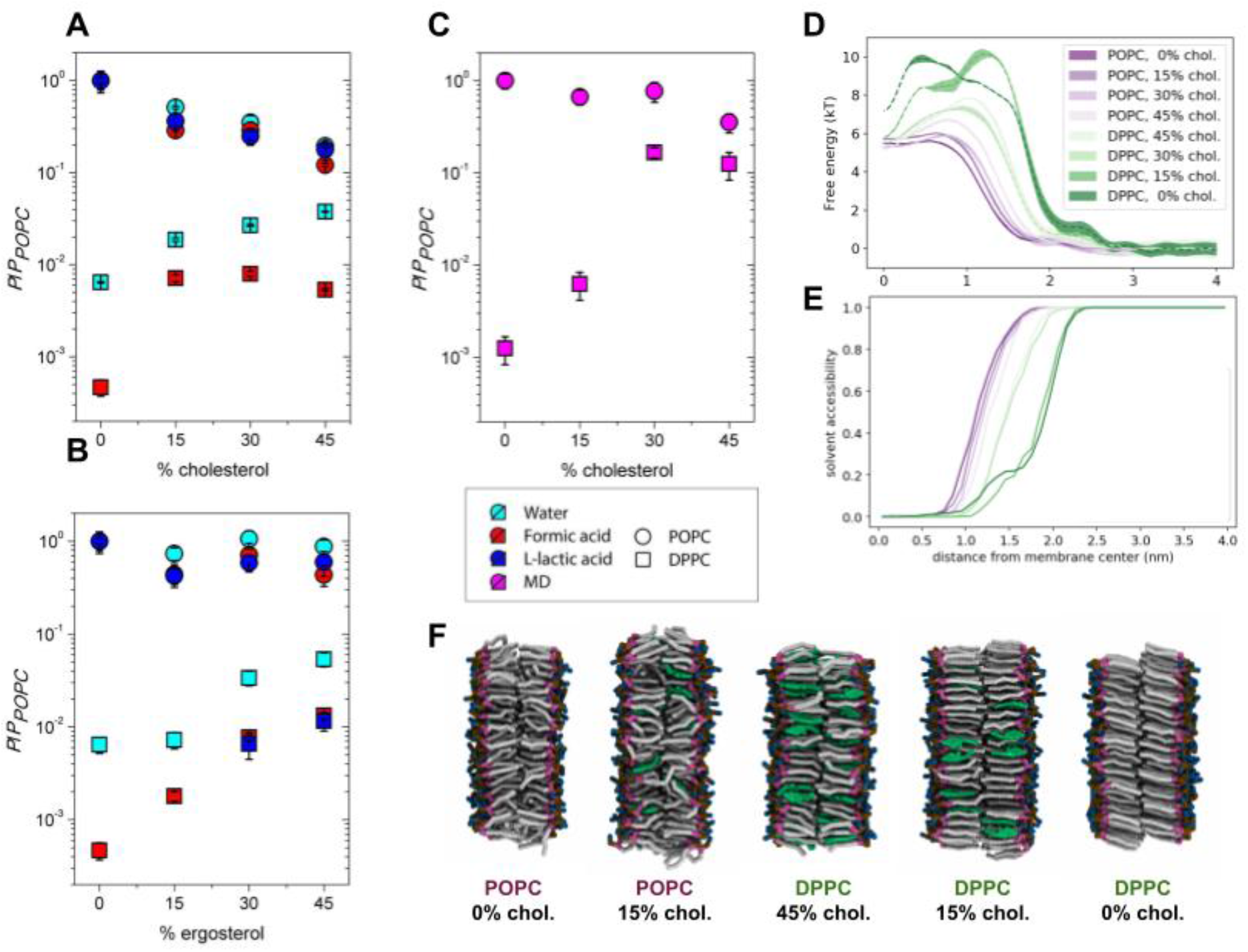
Permeability coefficients as a function of cholesterol (A, B) and ergosterol (C) concentration in POPC or DPPC vesicles from experiments (A, B) and simulations (C). The permeability coefficients for each solute are normalized to the value in POPC vesicles, *i.e*., without sterol present; the permeability coefficient in POPC vesicles was 12.0 (±1.4) x 10^-3^ cm/s, 5.53 ± (0.74) x 10^-3^ cm/s, and 0.117 (±0.022) x 10^-3^ cm/s for water, formic acid, and lactic acid, respectively. The numerical values are presented in Supplementary Table 4. D, E. Free energy (D) and solvent accessibility (E) profiles from simulated membranes in the L_d_ phase and L_β_ or L_O_ phase at varying cholesterol concentrations. The largest changes in the profiles and in the permeability coefficients are between simulations in different phase states. F. MD simulation snapshots of lipid membranes of various compositions at different phase states (lipid tails grey; glycerol moiety, purple; choline moiety, blue; phosphate moiety, ochre; cholesterol, green; water molecules are not shown).

For the membranes with POPC (*d*=0.5), both ergosterol and cholesterol lead to a small but significant decrease in permeability. In line with the known smaller condensing effect of ergosterol compared to cholesterol on lipid bilayer structure^29,30^, the decrease in permeability coefficient upon adding 45 mol% sterols is only approximately 2-fold for ergosterol but 5-fold for cholesterol. Similar effects are observed when titrating both sterols in vesicles composed of DOPC (Supplementary Table 4 and Supplementary Fig. 5). Our simulations show that in analogy to a decreasing unsaturation index in membranes without sterols (Fig. 3), the permeability coefficient decreases with increasing sterol concentration in POPC vesicles because of higher and wider free energy barriers (Fig. 4D). The effects of adding cholesterol are smaller in simulations than in the experiments but they are in line with the general trend.

The phase change from L_d_ to L_o_ in the simulations with POPC and between 15% and 30% sterol leads to a notable shoulder in the profile around 1.0 nm distance from the membrane center, where it is accompanied by a small depression in the profile (0.0 nm) (Fig. 4D). The corresponding changes of the permeability coefficients and free energy profiles are in accordance with our experiments (Fig. 4 and Supplementary Fig. 5) as well as with the recent simulations that compare the permeability of water through membranes in L_O_ and L_d_ phases^18^. The characteristic features of the free energy profiles of the L_O_ phase are linked to the sterol-induced changes in lipid packing^31^ and changes in the lateral pressure profile^32^. Unless a phase transition occurs, the shoulder in the profile continues to smoothly build up with decreasing unsaturation index towards DPPC (*i.e*., towards d=0.0; Supplementary Fig. 6).

Titration of sterols into fully saturated DPPC bilayers (*d*=0.0) has the opposite effect on solute permeability than in POPC membranes. For instance, we observe an overall increase of the permeability coefficient of about one order of magnitude between DPPC membranes in the absence or presence of 45 mol% of cholesterol or ergosterol. This large difference is caused by the change of the membrane phase from L_β_ to L_O_ upon addition of sterols. Opposite to what is seen in POPC membranes, sterols perturb the highly ordered acyl chains of the DPPC lipids leading to a significant decrease of the friction coefficient, *e.g*., compare DPPC bilayers to DPPC plus 15% cholesterol (Supplementary Fig. 3). As the free energy profiles for the DPPC membrane with 0 and 15 mol% of cholesterol are comparable in height and width, the change in the friction through the membranes in the L_β_ phase is an important factor for the difference in the permeability coefficient between the two lipid compositions.

Interestingly, the added cholesterol prevents water molecules from following the permeating solute into the membrane hydrophobic core. While water molecules can follow the permeating solutes into the DPPC membrane even below 1.0 nm (Fig. 4E), adding 15% cholesterol to the membrane keeps the water molecules below that threshold. This is linked to the shift of the free energy barrier peak from 0.5 nm in DPPC to 1.3 nm in DPPC plus 15% cholesterol and indicates that the sterols enforce solute dewetting further away from the hydrophobic region of the membrane. This effect comes from the highly anisotropic structure of the DPPC bilayer in the L_β_ phase, which forms “wedge-like” defects to accommodate the permeating solutes through its highly ordered structure and expose an accessible surface for the solvent molecules to enter “for free”; these cavities are filled with ergosterol or cholesterol when sterols are present in the membrane.

Adding more than 15% sterols to DPPC membranes leads to a change in the membrane phase from L_β_ to L_O_ and another order of magnitude increase in the permeability coefficient, which is a direct consequence of the dramatically lowered free energy barrier (Fig. 4D). Increasing cholesterol from 30% to 45% in membranes in the L_o_ phase leads to a decrease of the permeability coefficient, similar to what is observed for membranes with POPC. As the compositions with more than 30% cholesterol are in the same L_o_ phase, this effect can be attributed to the condensing effect of cholesterol on lipid bilayer structure^29,30^. Thus, the permeability coefficients decrease with the membrane fluidity in the order L_d_ > L_O_ > L_β_, each transition leading to an order of magnitude change in the permeation for water and weak acids.

### Differences in permeability of DPPC membranes with ergosterol and cholesterol reflect their phase behavior

The transitions between L_O_ and L_β_ phases differ for DPPC membranes with ergosterol and cholesterol. While 15 mol% of cholesterol in DPPC membranes increases the permeability coefficient for formic acid approximately 15-fold, the increase is only 5-fold for the same amount of ergosterol (Fig. 4B). Also, different from cholesterol, increasing the concentration of ergosterol in DPPC membranes up to 45% leads to a gradual increase of the permeability coefficient for formic acid rather than a more abrupt increase as seen between 0 and 15% cholesterol, suggesting a smooth change from L_β_ to L_O_ through phase coexistence.

A DPPC membrane with 15 or 30 mol% of cholesterol or ergosterol coexists of L_β_ and L_O_ phases (Supplementary Table 4) as confirmed by our DSC measurements (Supplementary Fig. 4B and C). The DPPC membrane with 15% of cholesterol forms a stable L_β_ phase in our MD simulations, but the DPPC membrane with 30% cholesterol forms a stable L_o_ phase, yielding a large change in the free energy profile and a corresponding increase in the permeability coefficient. Comparison of the changes of the permeability coefficients between simulations and experiments suggests that ergosterol has a higher tendency to form a L_β_ phase than cholesterol^33^, at the same concentration; cholesterol preferably forms a Lo phase. This is well in line with the observed overall smaller effects of ergosterol on the lipid bilayer structure compared to that of cholesterol^20,29,30^.

It may also explain the intriguing behavior of L-lactic acid, which has a much lower (almost 50 fold lower) permeability coefficient in POPC membranes than formic acid, while its permeation is not detectable in DPPC membranes w/o cholesterol (Fig. 4A and Supplementary Table 4). However, we did detect a measurable permeation of L-lactic acid in DPPC vesicles with 30% and 45% of ergosterol (Fig. 7B). This observation is in line with the known higher condensing effect of cholesterol compared to ergosterol and that cholesterol forms dimeric or even tetrameric aggregates at concentrations above 20%, which may be responsible for the observed permeation slowdown compared to ergosterol^29,30,34,35^.

## Discussion

We have used kinetic flux measurements and MD simulations to systematically analyze the permeability of small molecules of different polarity and size through membranes of varying lipid compositions. The used lipid compositions span different head groups, acyl chain lengths, degrees of unsaturation and the effects of cholesterol and ergosterol, allowing us to study the permeation through membranes in various phases, namely L_d_, L_O_ and L_β_.

Our MD simulations show that the partitioning of hydrophilic solutes to the membrane interior is comparable to their partitioning in octanol, but hydrophobic compounds partition into the membrane less than they do in octanol (Fig. 1). This highlights the limitations of using the octanol-water partition coefficient for estimating the permeability coefficients of hydrophobic compounds. These differences are reflected in the corresponding free energy and solvent proximity profiles (Fig. 1C and D). Unlike bulk solvents which are used to measure the partition coefficient such as *P_OW_*, membranes are comparably thin anisotropic layers. Hence, the process of solute permeation occurs for a large part through the mixed region at the interface including both the polar solvent and the hydrophobic lipid tails affecting the partition coefficient (Fig. 1D).

### How do membrane properties affect solute permeability?

The membrane properties and the phase behavior in particular have a great influence on the permeability coefficients for small polar molecules. We show that in the case of fluid membranes their thickness is the main parameter of the permeability coefficient (Fig. 2 and Supplementary Fig. 7). We find no general correlations between permeability coefficients and lipid surface areas contrary to what was observed by Mathai and colleagues^4^. Instead, we observe that adding carbon atoms to the lipid tails progressively increases the membrane hydrophobic thickness and decreases the permeability coefficients by an average of ca. 1.5 folds every two carbons in the range from 14 to 22 carbons.

We further demonstrate the dependence of the permeability coefficient on the membrane hydrophobic thickness also in membranes with different degrees of unsaturation and sterol content (Supplementary Figure S6). In contrast, lipid headgroup composition has only limited impact on the permeation of small molecules (Supplementary Figure S8 and Supplementary Table 5).

The relationship between permeability and membrane physical state (L_d_ and L_O_ phases) has received some attention in the past decade^18,36^. In line with these studies, we observe that the fluidity of the membranes decreases with decreasing degree of lipid tail unsaturation, and the rate of permeation decreases for both monounsaturated as well as polyunsaturated lipid bilayers^14,37^. The effect on solute permeability of changing the lipid unsaturation index (*d*) is relatively small for membranes in the fluid phases L_d_ and L_O_, and the difference in permeability coefficient between POPC (*d* = 0.5) and DOPC (*d* = 1.0) membranes is only about 5-fold. Changes of much larger magnitude appear with membrane phase transitions. We observe changes of three orders of magnitude for the permeability of formic acid through membranes in L_d_ and L_β_ phase, and between one and two orders of magnitude through membranes in L_O_ and L_β_ phases.

The exact values depend on the membrane composition and on the proximity to the phase transition temperature. We show in our simulations that the large differences in permeability arise from the different free energy profiles of the different membrane phases, which acquire small shoulders at the membrane-water interface when sterols are present, in line with existing studies^18^. In addition, the pure DPPC membrane (L_β_ phase) shows highly decreased diffusivity through its hydrophobic core (Supplementary Fig. S7). Adding sterols to the DPPC membrane without changing the membrane overall phase leads to an increase in the diffusivity towards the level of membranes in L_O_. Importantly, the sterols prevent the solutes to drag solvent molecules into the hydrophobic region of the membrane, thereby increasing the hydrophobic thickness (Fig. 4E).

Our measurements reveal important differences between cholesterol and ergosterol, which relate to the differences these sterols have on the phase behavior of the membrane. If different lipid phases coexist or the lipid bilayer is close to its phase transition temperature, the majority of solutes will diffuse in or out through the most permeable parts of membrane and/or the interphase with the coexisting phase^38^. Our results support the view that ergosterol has a higher tendency to form a L_β_ phase^33^, while cholesterol forms an L_O_ phase at the same concentration. This is in line with the observed overall smaller effects of ergosterol on the lipid bilayer structure compared to that of cholesterol^20,29,30^.

### How do our *in vitro* results relate to cell membranes *in vivo*?

The plasma membrane of yeast is laterally heterogeneous with a complex organization, specialized compartments and highly ordered rigid domains, which have inspired the here-presented measurements on the permeation of small hydrophilic solutes through membranes in different physical states (L_d_, L_O_ and L_β_). We and others have found that the lateral diffusion coefficient of proteins in the plasma membrane of yeast is 3-orders of magnitude slower than that of similar size proteins in the ER or vacuolar membrane^39–42^. The lateral diffusion of proteins in the plasma membrane of yeast is also much slower than in *e.g*. bacterial membranes^43^. Moreover, we have observed that the permeability of the yeast plasma membrane for formic acid and acetic acid is two to three orders of magnitude lower than that of synthetic lipid vesicles, and that the yeast membrane is virtually impermeable (at the level of passive diffusion) for lactic acid and glycerol, whereas bacterial membranes rapidly permeate these molecules^14^.

We have hypothesized that this difference in protein diffusivity and solute permeability is due to a more ordered state of the yeast plasma membrane. Specifically, the presence in the PM of long saturated acyl chain(s) sphingolipids, namely phosphoceramide (IPC), mannosyl-inositol-phosphoceramide (MIPC), mannosyl-(inositolphospho)2-ceramide (M(IP)2C), and ergosterol, particularly concentrated in the cytoplasmic leaflet^44^, could lead to a highly ordered structure. This hypothesis is in accordance with the work of Aresta-Branco and colleagues^19^, who studied plasma membrane of *Saccharomyces cerevisiae* by performing anisotropy measurements of diphenylhexatriene and fluorescent lifetime measurements of trans-parinaric acid. Estimates of global membrane order in wildtype cells and studies of mutants defective in sphingolipid or ergosterol synthesis suggest that the yeast plasma membrane harbors highly-ordered domains enriched in sphingolipids. In fact, the observed fluorescence lifetimes (>30 ns) are typical of the gel phase of synthetic membranes^45–48^.

In recent work^49^, we observed a high degree of unsaturated acyl chains and low values of ergosterol in the very small shell of lipids surrounding membrane transport proteins. These proteins with the so-called periprotein lipidome are embedded in an environment of lipids that are enriched in ergosterol and possibly saturated long-chain fatty acids such as present in IPC, MIPC and M(IP)_2_C), which yield a highly liquid-ordered state. The highly ordered state likely explains the slow lateral diffusion and the low solute permeability of the yeast plasma membrane. The ordered state may also form the basis for the robustness of yeast to strive in environments of low pH, high concentrations of ethanol or weak acids, and relate to the ability of *S. cerevisiae* cells to retain glycerol and use it as main osmoprotectant^50–55^.

We now show that the permeability coefficients for formic acid, lactic acid, and water^56^ in DPPC DPPC/ergosterol and DPPC/cholesterol vesicles are in line with the observations on permeation and lateral diffusion made for the yeast plasma membrane (Table 1). We were not able to test the long saturated acyl chain(s) sphingolipids of yeast, because IPC, MIPC and M(IP)2C are not available and we used DPPC instead. The permeability of the yeast plasma membrane for water and formic acid is at least two orders of magnitude slower than of vesicles in the liquid disordered phase, *i.e*., DOPC vesicles. On the other hand, membranes in the gel and liquid-ordered phase, *i.e*., DPPC vesicles with or without 15% ergosterol or cholesterol, display permeability coefficients similar to that of the yeast plasma membrane. We also notice that the water/formic acid and water/glycerol permeability ratio follow the order L_d_ < L_O_ < L_β_’ (*e.g*., DOPC < DPPC + 30% sterol < DPPC). This indicates that formic acid and glycerol permeation through ordered bilayers is more penalized compared to water diffusion. Interestingly, in yeast we observe a water/formic acid ratio that lays between the ones of pure DPPC and DPPC + 30% cholesterol. We have also shown that yeast PM is impermeable to lactic acid in the timespan of 2.5 hours. To our knowledge, passive influx of lactic acid has not been observed in *S. cerevisiae* cells, which parallels the observation that we do not observe lactic acid permeation in vesicles composed of DPPC or DPPC:cholesterol. Yet, we observe slow permeation in vesicles of DPPC:ergosterol (=70:30, 55:45), where the Lo phase predominates. The evidence collectively corroborates that the yeast plasma membrane behaves as a highly ordered barrier to the permeation of small polar molecules, which is different from the more fluid plasma membranes of mammalian and prokaryotic cells.

**Table 1.** Permeability coefficients in cm/s of water, formic acids, lactic acid and glycerol for lipid vesicles in different physical states and yeast cells. *Permeability coefficient for water in double mutant *aqy1 aqy2*^56^; **permeability coefficient for formic acid in RA380^14^.

| | $P_{DOPC}$<br>$\times 10^{-5}$ (cm/s) | $P_{DPPC}$<br>$\times 10^{-5}$ (cm/s) | $P_{DPPC+15\%cholesterol}$<br>$\times 10^{-5}$ (cm/s) | $P_{DPPC+15\%ergosterol}$<br>$\times 10^{-5}$ (cm/s) | $P_{yeast}$<br>$\times 10^{-5}$ (cm/s) |
| --- | --- | --- | --- | --- | --- |
| Water | 1600 ± 170 | 7.8 ± 1.0 | 23 ± 4 | 12 ± 2 | 10-20* |
| Formic acid | 719 ± 172 | 0.27 ± 0.05 | 4.1 ± 0.3 | 1.3 ± 0.1 | 1.1 ± 0.1** |
| L-lactic acid | 19.8 ± 3.3 | Not observed | Not observed | Not observed | Not observed |
| Glycerol | 0.23 ± 0.02 | 0.00011 ± 0.00002 | 0.001284 ± 0.00013 | / | - |

Our data are in line with the suggestion that in yeast the liquid-disordered phase is confined to the periprotein lipids^49^. The high degree of unsaturated acyl chains and low values of ergosterol in the periprotein lipidomes may allow sufficient conformational flexibility of the proteins, yet without compromising the exceptional permeability barrier of the membrane. The high degree of saturated lipid in the yeast plasma membrane^57^ compared to for instance the membranes of human cells may be seen as a possible evolutionary adaptation of yeasts to ergosterol as their main sterol. It is clear from the differences between ergosterol and cholesterol, that the two sterols exhibit different phase behavior and have different interactions with saturated and unsaturated lipid tails^20,29,30,33^. In agreement, our measurements with saturated DPPC lipids show that cholesterol exhibits large jumps in permeability, fluidity and, hence, in phase behavior, whereas ergosterol impacts the membrane properties and phase state more smoothly. This allows ergosterol to function as a component that can steadily regulate fluidity in the highly rigid yeast plasma membranes but less so in more unsaturated fluid membranes, where cholesterol is a better regulator.

## Conclusions

In summary, we find that the membrane thickness and the degree of lipid tail unsaturation have a significant impact on the solute permeability in membranes in the fluid phase, but the biggest changes in permeation happen when these factors lead to the transition from the fluid (L_d_) to the gel-like phase (L_β_). We observe a drop of three orders of magnitude in the permeability of formic acid and glycerol, and two orders of magnitude in water permeability for DPPC, compared to DOPC membranes. This is mainly due to the large differences in the solubility of the permeating solutes in the membrane interior. The addition of cholesterol or ergosterol to DPPC, which coincides with the formation of the liquid-ordered phase (L_O_), induces a partial restoration of permeability towards the level of fluid membranes. Our measurements reveal that ergosterol has a small impact on lipid bilayer structure compared to cholesterol, with the latter having a higher tendency to induce a L_O_ phase at the same concentrations. We also find that, when more phases coexist, solutes preferentially permeate through the more fluid parts of the membrane and/or through the region between the coexisting phases. Finally, we compare our results with *in vivo* data from *Saccharomyces cerevisiae* presented previously and use the permeability data as reporter of the physical state of the yeast plasma membrane. Our results reveal that the yeast plasma membrane is in a highly rigid physical state comparable to model membranes of DPPC with 0–15% ergosterol at a gel-like L_β_ phase. Moreover, unlike cholesterol, ergosterol changes the membrane properties including the permeability coefficient smoothly with its concentration in that regime allowing it to act as a membrane rigidity regulator in yeasts.

## Materials and methods

### Materials

The weak acid solutions were prepared using the following salts: sodium-acetate (BioUltra, ≥99.0%; Sigma-Aldrich); sodium-benzoate (BioXtra, ≥99.5%, B3420-250G; Sigma-Aldrich, St. Louis, MO); sodium-butyrate (≥98.5%; Sigma-Aldrich); sodium-formate (BioUltra, ≥99.0%; Sigma-Aldrich); sodium L-lactate (>99.0%; Sigma-Aldrich); sodium-propionate (≥99.0%; Sigma-Aldrich); pyruvic acid-sodium salt (99+%; Acros Organics, Geel, Belgium); potassium-sorbate (purum p.a., ≥99.0%; Sigma-Aldrich): and potassium chloride (pro analyses; BOOM Laboratorium Leveranciers, Meppel, The Netherlands). Glycerol was purchased from BOOM Laboratorium Leveranciers (Meppel, The Netherlands). The following lipids were used and purchased fromAvanti Polar Lipids (Alabaster, AL): 1,2-dioleoyl-*sn*-glycero-3-phosphocholine (DOPC); 2-dioleoyl-*sn*-glycero-3-phosphoethanolamine (DOPE); 1,2-dioleoyl-*sn*-glycero-3-phospho-(1’-rac-glycerol) sodium salt (DOPG); 1-palmitoyl-2-oleoyl-*sn*-glycero-3-phosphocholine (POPC); 1-palmitoyl-2-oleoyl-*sn*-glycero-3-phosphoethanolamine (POPE); 1-palmitoyl-2-oleoyl-*sn*-glycero-3-phospho-(1’-rac-glycerol) sodium-salt (POPG); 1,2-dipalmitoyl-*sn*-glycero-3-phosphocholine (DPPC); 1,2-dimyristoleoyl-*sn*-glycero-3-phosphocholine (14:1 (Cis) PC), 1,2-dipalmitoleoyl-*sn*-glycero-3-phosphocholine (16:1 (Δ9-Cis) PC), 1,2-dieicosenoyl-*sn*-glycero-3-phosphocholine (20:1 (Cis) PC); 1,2-dierucoyl-*sn*-glycero-3-phosphocholine (22:1 (Cis) PC); cholesterol; ergosterol.

### Weak acid solutions

The 1 M stock solutions (0.5 M for benzoic acid) were prepared by dissolving the salt, or glycerol, into 100 mM potassium phosphate (KPi) and the pH was adjusted to 7.0 using 4 M NaOH. An empirical linear relation (y = mx + q) between osmolyte concentration and osmolality was determined for each solution (Supplementary Fig. 9). The osmolality was measured using a freezing point depression osmometer (Osmomat 3000 basic; Genotec, Berlin, Germany). The empirical relations were used to estimate the osmolyte concentrations needed for an osmolality of ~300 mosmol/kg, that is, upon mixing with the liposome solution. The stock solutions were accordingly diluted to the desired concentration before the experiment.

### Vesicle preparation

The lipids were purchased from Avanti Polar Lipids in powder and suspended in chloroform to a concentration of 25 mg/mL. After mixing the solubilized lipids in the desired ratio, a rotary vaporizer (rotavapor r-3 BUCHI, Flawil, Switzerland) was used to remove chloroform by evaporation. Next, the lipids were suspended in diethylether and subjected to a second of evaporation. Finally, the lipids were hydrated in the assay buffer (100 mM KPi, pH 7) and adjusted to a concentration of 10 mg/mL. The lipid solution was homogenized by tip (3.18 mm) sonication with a Sonics Vibra Cell sonicator (Sonics & Materials Inc. Newtown, CT, USA) at 4 °C (ice water) for 4 min with 15 sec pulses and 15 sec pause between every pulse. Amplitude of the sonicator was set to 100%. The prepared vesicles were stocked at 20 mg/mL in liquid nitrogen to prevent oxidation.

### Definition of the degree of unsaturation

The degree of unsaturation, *d*, equals the ratio between the number of lipid tails with carbon-to-carbon double bonds (*N_C=C_*) and the total number of tails (*N_total_*): *d* = *N_C=C_*/*N_total_*. All the phospholipids used in this work have two tails per head group and utmost one double bond per tail. Pure mixtures of DOPC, POPC and DPPC have degrees of unsaturation of 1, 0.5, and 0, respectively. Intermediate values of *d* were obtained by mixing DOPC and POPC (i.e., *d* between 1.0 and 0.5) or POPC and DPPC (i.e., *d* between 0.5 and 0.0).

### Preparation of vesicles filled with calcein

The fluorophore calcein (from Sigma-Aldrich) was solubilized at a concentration of 100 mM with 50 mM KPi, and the pH was adjusted to 7.0 using aliquots of 4 M KOH. The stocked vesicles (2 mg of lipid) were pelleted by ultracentrifugation (80,000 rpm, 4°C, 20 min with a TLA 100.1 rotor in a Beckman Optima TLX Ultracentrifuge; Beckman Coulter Life Sciences, Indianapolis, IN) and resuspended in 0.9 mL of 89 mM KPi, pH 7.0. Calcein was added to the liposome solution at a self-quenching concentration (10 mM) and enclosed in the vesicles by 3 cycles of rapid freezing in liquid nitrogen and thawing at 40°C (or 60°C for mixtures containing DPPC). Thus, the osmolality of the liposome lumen (filled with 10 mM calcein plus 89 mM KPi pH 7.0) is ~190 mosmol/kg, which equals the osmolality of the assay buffer (100 mM KPi pH 7.0). After extrusion through a 200 nm polycarbonate filter at 20°C (or 60°C for mixtures containing DPPC) to homogenize the vesicles, they were eluted through a 22-cm-long Sephadex-G75 (Sigma-Aldrich) column pre-equilibrated with the assay buffer to remove the external calcein. The collected 1 mL fractions containing the calcein-filled vesicles were identified by eye using an ultraviolet lamp (for fluorophore excitation) and diluted in a total volume of 10 mL of the assay buffer.

### Stopped-flow experiments

A stopped-flow apparatus (SX20; Applied Photophysics, Leatherhead, Surrey, UK) operated in single-mixing mode was used to measure fluorescence intensity kinetics upon application of an osmotic shock to the vesicles filled with calcein. To impose the osmotic shock, the solution of the permeant, weak acid in most cases (ca. 100 mM of sodium or potassium salt of the weak acid in 100 mM KPi pH 7.0; ~300 mosmol/kg after mixing), and the vesicles were loaded each in one syringe and forced first through the mixer (1:1 mixing ratio with 2 ms dead time) into the optical cell (20 μL volume and 2 mm pathlength). The temperature of the optical cell was set at 20°C using a water bath. The white light emitted by a xenon arc lamp (150 W) was passed through a high-precision monochromator and directed to the optical cell via an optical fiber. The band pass of the monochromator was optimized and set to 0.5 nm (for calcein) to prevent fluorophore photobleaching during the experiment. Calcein was excited at 495 nm. The emitted light, collected at 90°, was filtered by a Schott long-pass filter (cutoff wavelength at 515 nm) and detected by a photomultiplier tube (R6095; Hamamatsu, Hamamatsu City, Japan) with 10 μs time resolution. The voltage of the photomultiplier was automatically selected and kept constant during each set of experiments. The fluorescence intensity kinetics after the osmotic shock was recorded with logarithmically spaced time points to better resolve faster processes. For noise reduction, multiple acquisitions (three for slow kinetics and nine for fast kinetics) were performed for each experimental condition.

### Preprocessing of the in vitro kinetic data

The raw data were preprocessed in MATLAB (R2020b; The MathWorks, Natick, MA) for further analysis. First, the N number of curves, which we called fi(t), acquired with a single experimental condition, were averaged (F(t) = N–1∑fi(t)) to reduce the noise. For calcein, the resulting kinetic curves F(t) were normalized to 1 at time zero (F(t)/F(0)), i.e., the mixer dead time (t_0_ = 2 ms).

### Size distribution of vesicles

The size distribution of vesicles was measured by Dynamic Light Scattering (DLS) using the DynaPro NanoStar Detector (Wyatt Technology, Santa Barbara, CA). Empty vesicles were prepared starting from 1 mg of lipids by three freeze-and-thaw cycles at 40°C (or 60°C for mixtures containing DPPC). After 13 times extrusion through a 200 nm filter, vesicles were eluted through a 22-cm-long Sephadex-G75 column pre-equilibrated with 100 mM KPi (pH 7.0). Before the DLS measurements, the vesicles were diluted with the assay buffer to a concentration in the range from 2 μg/mL to 2 mg/mL. Measurements were performed with a scattering angle of 90°. For each measurement, at least 10 acquisitions of 20 s each were performed at a temperature of 20°C. For each acquisition, at least 2 million counts were recorded. The correlation curves and the intensity-weighted distributions were obtained with the built-in analysis software.

### Fit of the in vitro kinetics

The function <F(t)>/<F(0)> describes the time evolution of the calcein fluorescence and is calculated as in work^15^. Briefly, the relaxation kinetics of the calcein concentration c2(r_0,t_) was computed by numerical solution of the system of differential equations describing the dynamics of a spherical vesicle of radius r_0_ upon osmotic upshift. The numerical solution was used to calculate the ratio F(r_0,t_)/F(0), using the Stern-Volmer equation with dynamic quenching constant K_SV_. The population-averaged ratio <F(t)>/<F(0)> was computed by using the vesicle size distribution gi(r_0_) measured in dynamic light scattering (DLS, Supplementary Fig. 10) experiments and fitted in MATLAB to the experimental data using the FMINUIT^59^ minimization routine. For the “impermeable” osmolyte (KCl), two fitting parameters were used: the quenching constant K_SV_ (M^−1^) and the water permeability coefficient P_w_ (cm/s). For the permeable osmolytes (Table 2), the water permeability coefficient P_w_ was fixed to the value obtained from the KCl data, whereas K_SV_ and the permeability coefficient P_AH_ were fitted to each other. To improve the accuracy and to estimate the error of P_AH_, we repeated the fit for each of the 10 size distributions acquired by DLS. The mean of the fitted values was used as the best estimate of P_AH_, and the standard deviation indicates the experimental uncertainty for the permeability coefficient δP_AH_, which derives mostly from the ambiguity of the vesicle size distribution estimation by DLS. The other parameters required for calculation of c2(r_0,t_) were set to their experimental values, which are pH_out_ = 7, [KPi]_in_ = 89 mM, [KPi]_out_ = 100 mM, c2(r_0,0_) = 10 mM, M_W_ H_2_O = 18 cm^3^/mol, pK_a_ (KPi) = 7.21, and see Table 2 for pK_a_ (acid). M_W_ H_2_O is the molar volume of water. The concentration of the weak acids [AH]_out_ in the external solution was set to 40-50 mM for all compounds, except for glycerol, which was set to 120 mM, as obtained from Supplementary Fig. S9. Accordingly, the total osmolyte concentration is 2[AH]_out_ to account for the counterion released by the weak acid salt.

**Table 2.** Molecular Weight and pK_a_ values of the used osmolytes. The pKa values were taken from the PubChem database^58^, using the compound ID indicated in the last column. N/A, not applicable.

| Osmolyte | M<sub>w</sub> (g/mol) | pK<sub>a</sub> (25°C) | Compound ID |
| --- | --- | --- | --- |
| KCl | 74.55 | N/A |  |
| Sodium acetate | 82.03 | 4.76 | 176 |
| Sodium benzoate | 144.1 | 4.19 | 243 |
| Sodium butyrate | 110.09 | 4.82 | 264 |
| Sodium formate | 68.01 | 3.75 | 284 |
| Sodium L-lactate | 112.06 | 3.86 | 612 |
| Sodium propionate | 96.06 | 4.88 | 1032 |
| Sodium pyruvate | 110 | 2.45 | 1060 |
| Potassium sorbate | 150.22 | 4.76 | 643460 |
| Glycerol | 92.09 | 14.4 | 753 |

### Differential scanning calorimetry (DSC)

Differential scanning calorimetry (DSC) measurements were performed using an MC-2 calorimeter (MicroCal, Amherst, MA) to perform ascending and descending temperature mode operations. The lipid concentration used was 1–2 mg/mL and the temperature of the sample and reference cells was controlled by a circulating water bath. The scan rate was 20°C/h for both heating and cooling scans. Data were analyzed using ORIGIN software provided by MicroCal. Samples were scanned 5 times to ensure the reproducibility of the endotherms.

### Molecular dynamics (MD) simulations

MD simulation is an established method to study permeability of solutes through lipid membranes^9,16–18^. Here we rely on the latest version of the CG Martini model^25^. Bereau and co-workers have shown that the Martini model is very accurate for computing permeation rates, with good correlations to both all-atom simulations and experimental measurements over a wide range of compounds^23,24^. In the current work, permeants are modeled as generic single-particle solvents of varying hydrophobicity from I (most hydrophilic) to IX (most hydrophobic). The initial conditions of the simulated lipid bilayers were generated using the tools Insane and Martinate^60^ to yield lateral dimensions of 10×10 nm with the bilayer repeat distance of 10 nm. The systems were equilibrated at their respective temperatures for at least 10 ns prior to production simulations. A standard simulation setup for the Martini model was used as described in previous work^25^. In brief, the simulation temperature was coupled to a v-rescale thermostat^61^ at room temperature of 293 K, separately for the lipids and solvent. A Parinello-Rahman barostat^62^ was used for pressure coupling at 1 bar with a coupling constant of 24 ps independently for the membrane plane and its normal. Standard time step of 20 fs was used for all simulations and the trajectory was recorded every 1 ns. For production, we have simulated all systems for 5 μs to ensure convergence; simulations of membranes at the gel phase were run for 10 μs. Simulations were run using GROMACS simulation package ver. 2019.3 in a mixed precision compilation without GPU support^62,63^. Adaptive weighted histogram (AWH) method^64^ was used to obtain free energy profiles of the permeating particles. The AWH weight factors were updated every 0.1 ns. Particle permeation was done along the z-axis as a membrane normal. Distance of the permeating particle from the membrane center was calculated using the cylindrical coordinate, which uses the local membrane lipids within a cutoff distance of 1.0 nm to calculate the center of mass of the membrane^63^. The force constant of the biasing potentials in AWH was 2000 kJ·mol^-1^·nm^-2^ and the coupling constant between the generalized and the molecular coordinates was 10000 kJ·mol^-1^·nm^-2^. The constant of local friction was calculated from force autocorrelation implemented within the AWH method^63,64^. The friction profiles were denoised using a Butterworth lowpass filter^65^ of the fourth order with a Nyquist frequency of 100 Hz. Error estimate was calculated from the average noise around the denoised curve and from the deviations from the symmetrical shape between the left and right part of the profile. The profiles of solvent proximity to the permeants were calculated using GROMACS tool gmx mindist with a cutoff radius of 0.6 nm between the permeant and solvent molecules. Scripts used to generate and analyze the simulations are available in an open public repository^66^.

### Inhomogeneous solubility-diffusion model

The inhomogeneous solubility-diffusion (ISD) model was used to calculate the permeability coefficient *P* using Equation 1

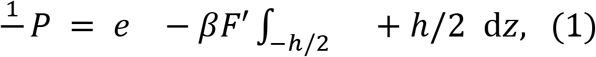

where *F(z)* denotes the free energy as a function of the membrane normal, *D(z)* denotes the local diffusivity, *h* is the thickness of the membrane, and □ = 1/(*k_B_T*) with the Boltzmann constant *k_B_* and temperature *T*. The exponential term in Equation 1 represents the local solubility as a function of the membrane normal, *K(z)*, calculated from the free energy, *F(z)*. Local friction, *f(z)*, is inversely proportional to the local diffusivity as *f(z)* = *kT*/*D*(*z*). Error estimates were calculated using standard chain rules for propagating errors, which arise mainly from the uncertainty of the free energy profiles.

The permeation dynamics is assumed to be diffusive in the ISD model. Deviations from such an assumption may lead to inaccurate estimates of the permeability coefficients for permeants that interact strongly with one another or with the membrane^67,68^. Also, for higher concentrations, the ISD model does not capture the collective behavior of the permeating solute and its effects on the membrane. However, for compounds at low concentrations the estimates of the permeability coefficients from the ISD model are comparable to the unbiased estimates from transition-based counting^68^. Further studies on the permeability of small solutes through lipid membranes and the ISD in various scenarios are provided in the references^16,67–70^.

By assuming average behavior over the permeation pathway in Equation 1, we obtain a simpler homogeneous solubility diffusion (HSD) model, also known as the Meyer-Overton rule

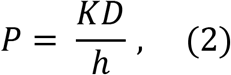

with *h* denoting the membrane hydrophobic thickness. Other symbols have the same meaning as above in Equation 1.

### Calculation of the solvent accessibility profiles, partition coefficients and the membrane hydrophobic thickness

Solvent accessibility profiles represent the local probability of the permeating molecule to be in a contact (within cutoff distance of 0.6 nm) with at least one solvent molecule at the given distance from the membrane center.

Membrane hydrophobic thickness, *h*, was calculated after the definition of Luzzati^71^ according to the formula

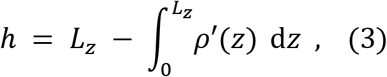

where *L_z_* is the simulation box length is *z*-direction (also known as bilayer repeat distance), and □’*(z)* is the local density of the solvent normalized to the value in the bulk.

The partition coefficients were obtained directly from the free energy profiles as the difference

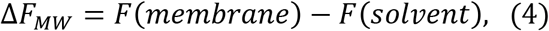

where *F*(*z*) is the calculated free energy at the membrane center and in the solvent bulk phase, respectively. The octanol-water partition coefficients were taken directly from^25^.

## Acknowledgements

We thank Matteo Gabba for insightful discussions at the start of the project. The research was funded by an ERC Advanced grant (ABCVolume; #670578) and the EU CoFund program ALERT. We thank SURFsara and the Center for Information Technology of the University of Groningen for their support and for providing access to the high-performance computing clusters Cartesius and Peregrine.

