## Supplementary data for "Membrane thickness, lipid phase and sterol type are determining factors in the permeability of membranes to small solutes"

### Supplementary Tables

| Compound | Log$P_{OW}$ | Experimental permeability coefficients x$10^{-3}$ (cm $s^{-1}$) |
|---|---|---|
| Water | - | 16.0 ± 1.7 |
| Glycerol | -1.76 | 0.0023 ± 0.002 |
| L-lactic acid | -0.6 | 0.198 ± 0.033 |
| Formic acid | -0.54 | 6.73 ± 0.74 |
| Pyruvic acid | -0.34 | 1.20 ± 0.17 |
| Acetic acid | -0.17 | 95.1 ± 1.1 |
| Propionic acid | 0.33 | 101 ± 3 |
| Butyric acid | 0.79 | 214 ± 34 |
| Sorbic acid | 1.33 | 291 ± 30 |
| Benzoic acid | 1.87 | 336 ± 51 |

| Hydrophobicity level and simulated particle name | Log$P_{OW}$ | Log$P_{MW}$ | Calculated permeability coefficients x$10^{-3}$ (cm $s^{-1}$) |
|---|---|---|---|
| I, SP6 | -2.14 | -2.03 | 0.17 ± 0.07 |
| II, SP3 | -1.35 | -1.15 | 35 ± 2 |
| III, SP1 | -0.91 | -0.88 | 96 ± 11 |
| IV, SN5 | -0.63 | -0.66 | 402 ± 80 |
| V, SN3 | -0.32 | -0.33 | 2050 ± 200 |
| VI, SN2 | 0.37 | -0.06 | 7700 ± 800 |
| VII, SN1 | 0.63 | 0.11 | 10400 ± 1500 |
| VIII, SC6 | 0.93 | 0.26 | 12800 ± 2000 |
| IX, SC5 | 1.1 | 0.46 | 11600 ± 2000 |

**Supplementary Table 1**. Permeability coefficients in lipid vesicles composed of pure DOPC and octanol/water, and membrane/water partitioning coefficient (log$P_{OW}$, log$P_{MW}$) from experiments and simulations.

| Lipids | Acyl chain length | $P_{Water}$ x10$^{-3}$ (cm/s) | $P_{formic\ acid}$ x10$^{-3}$ (cm/s) | $P_{lactic\ acid}$ x10$^{-3}$ (cm/s) | $P_{MD}$ x10$^{-3}$ (cm/s) |
|---|---|---|---|---|---|
| (14:1) PC | 14 | 22.8 ± 1.9 | 11.80 ± 1.43 | 0.332 ± 0.052 | 199 ± 30 |
| (16:1) PC | 16 | 16.3 ± 1.1 | 8.31 ± 0.64 | 0.196 ± 0.012 | / |
| (18:1) PC/DOPC | 18 | 16.0 ± 1.7 | 7.19 ± 1.72 | 0.198 ± 0.033 | 96 ± 11 |
| (20:1) PC | 20 | 7.1 ± 0.9 | 3.39 ± 0.13 | 0.098 ± 0.011 | / |
| (22:1) PC | 22 | 5.9 ± 0.6 | 1.90 ± 0.22 | 0.078 ± 0.010 | 42 ± 2 |
| (26:1) PC | 26 | / | / | / | 28 ± 2 |

**Supplementary Table 2.** Permeability coefficients for formic acid, lactic acid, water, and the simulated solute of hydrophobicity level III (SP1) in lipid vesicles composed of pure PC with varying acyl chain length from experiments and simulations.

| Mixture | d | Expected phase | $P_{water}$ x10$^{-3}$ (cm/s) | $P_{formic\ acid}$ x10$^{-3}$ (cm/s) | $P_{lactic\ acid}$ x10$^{-3}$ (cm/s) | $P_{glycerol}$ x10$^{-6}$ (cm/s) | $P_{MD}$ x10$^{-3}$ (cm/s) |
|---|---|---|---|---|---|---|---|
| DOPC | 1 | $L_d$ | 16.0 ± 1.7 | 6.73 ± 0.74 | 0.198 ± 0.033 | 2.3 ± 0.2 | 96 ± 11 |
| DOPC/POPC | 0.84 | $L_d$ | 13.5 ± 0.5 | 6.04 ± 0.36 | 0.177 ± 0.021 | / | / |
| POPC/DOPC | 0.67 | $L_d$ | 12.3 ± 1.5 | 5.68 ± 0.48 | 0.153 ± 0.031 | / | / |
| POPC | 0.5 | $L_d$ | 12.0 ± 1.4 | 5.53 ± 0.74 | 0.117 ± 0.022 | 1.27 ± 0.1 | 48 ± 9 |
| POPC/DPPC | 0.34 | *$L_d + L_\beta$ | 8.7 ± 1.0 | 4.90 ± 0.92 | 0.083 ± 0.008 | / | 33 ± 7 |
| DPPC/POPC | 0.17 | *$L_d + L_\beta$ | 5.9 ± 0.3 | 2.36 ± 0.56 | 0.049 ± 0.004 | / | 31 – 0.08 |
| DPPC | 0 | * $L_\beta$ | 0.078 ± 0.010 | 0.0027 ± 0.0005 | Not observed | 0.0011 ± 0.0002 | 0.06 ± 0.02 |

**Supplementary Table 3.** Permeability coefficients for various compounds in lipid vesicles composed of pure DOPC, POPC and DPPC, and mixtures of DOPC/POPC and POPC/DPPC. Parameter *d* represents the degree of unsaturation. The errors (SD) originate from the inaccuracy of the vesicle size distribution. *Mixtures analyzed by DSC (Supplementary Fig. 4A).

| Lipid mixture | Expected phase | $P_{water}$ x$10^{-3}$ (cm/s) | $P_{formic\ acid}$ x$10^{-3}$ cm/s) | $P_{lactic\ acid}$ x$10^{-3}$ (cm/s) | $P_{MD}$ x$10^{-3}$ (cm/s) |
|---|---|---|---|---|---|
| DOPC | $L_d$ | 16.0 ± 1.7 | 6.73 ± 0.74 | 0.198 ± 0.033 | 96 ± 11 |
| DOPC + 15 % Chol | $L_d$ | 10.3 ± 1.1 | 3.10 ± 0.12 | 0.090 ± 0.012 | / |
| DOPC + 30 % Chol | $L_O$ | 7.69 ± 0.41 | 3.49 ± 0.35 | 0.047 ± 0.004 | / |
| DOPC + 45 % Chol | $L_O$ | 4.41 ± 0.47 | 0.58 ± 0.04 | 0.015 ± 0.002 | / |
| DOPC + 15 % Erg | $L_d$ | 10.1 ± 0.4 | 5.64 ± 0.26 | 0.076 ± 0.002 | / |
| DOPC + 30 % Erg | $L_O$ | 10.6 ± 1.1 | 2.45 ± 0.13 | 0.149 ± 0.012 | / |
| DOPC + 45 % Erg | $L_O$ | 12.1 ± 1.4 | 2.79 ± 0.11 | 0.150 ± 0.022 | / |
| POPC | $L_d$ | 12.0 ± 1.4 | 5.53 ± 0.74 | 0.117 ± 0.022 | 48 ± 9 |
| POPC + 15 % Chol | $L_d$ | 6.22 ± 0.34 | 1.65 ± 0.49 | 0.042 ± 0.002 | 32 ± 6 |
| POPC + 30 % Chol | $L_d$ + $L_O$ | 4.28 ± 0.70 | 1.67± 0.36 | 0.029 ± 0.002 | 37 ± 9 |
| POPC + 45 % Chol | $L_O$ | 2.38 ± 0.36 | 0.70 ± 0.02 | 0.021 ± 0.003 | 17 ± 3 |
| POPC + 15 % Erg | $L_d$ | 8.97 ± 0.40 | 2.54 ± 0.69 | 0.049 ± 0.003 | / |
| POPC + 30 % Erg | $L_d$ + $L_O$ | 12.9 ± 2.2 | 4.11 ± 1.38 | 0.068 ± 0.005 | / |
| POPC + 45 % Erg | $L_O$ | 10.7 ± 1.0 | 2.53 ± 0.62 | 0. 070 ± 0.002 | / |
| DPPC | *$L_\beta$ | 0.078 ± 0.010 | 0.0027 ± 0.0005 | Not observed | 0.06 ± 0.02 |
| DPPC + 15 % Chol | *$L_\beta$ + $L_O$ | 0.23 ± 0.04 | 0.041 ± 0.003 | Not observed | 0.3 ± 0.1 |
| DPPC + 30 % Chol | *$L_\beta$ + $L_O$ | 0.33 ± 0.02 | 0.046 ± 0.003 | Not observed | 8 ± 1 |
| DPPC + 45 % Chol | *$L_O$ | 0.46 ± 0.07 | 0.031 ± 0.008 | Not observed | 6 ± 2 |
| DPPC + 15 % Erg | * $L_\beta$ + $L_O$ | 0.12 ± 0.02 | 0.013 ± 0.001 | Not observed | / |
| DPPC + 30 % Erg | * $L_\beta$ + $L_O$ | 0.54 ± 0. 05 | 0.056 ± 0.003 | 0.0013 ± 0.0003 | / |
| DPPC + 45% Erg | *$L_O$ | 0.86 ± 0.07 | 0.096 ± 0.006 | 0.0023 ± 0.0003 | / |

**Supplementary Table 4.** Permeability coefficients for various compounds in lipid vesicles composed of pure DOPC, POPC and DPPC with cholesterol or ergosterol. *Mixtures analyzed by DSC (Supplementary Fig. 4B and C).

| Lipid mixture | Ratio | $P_{water}$ x$10^{-3}$ (cm/s) | $P_{formic\ acid}$ x$10^{-3}$ cm/s) | $P_{lactic\ acid}$ x$10^{-3}$ (cm/s) |
|---|---|---|---|---|
| DOPE:DOPC:DOPG | 0:75:25 | 13.0 ± 1.2 | 7.47 ± 0.83 | 0.119 ± 0.024 |
| DOPE:DOPC:DOPG | 25:50:25 | 13.2 ± 1.3 | 8.18 ± 0.75 | 0.151 ± 0.020 |
| DOPE:DOPC:DOPG | 50:25:25 | 15.3 ± 1.4 | 7.38 ± 0.93 | 0.121 ± 0.040 |
| DOPE:DOPC:DOPG | 60:15:25 | 9.2 ± 0.9 | 2.42 ± 0.13 | 0.069 ± 0.005 |
| DOPE:DOPC:DOPG | 70:05:25 | 5.1 ± 1.5 | 1.48 ± .015 | 0.047 ± 0.009 |
| DOPE:DOPC:DOPG | 50:50:0 | 13.8 ± 1.9 | 6.37 ± 1.09 | 0.116 ± 0.016 |
| DOPE:DOPC:DOPG | 50:37:13 | 13.2 ± 1.4 | 8.26 ± 1.08 | 0.117 ± 0.016 |
| DOPE:DOPC:DOPG | 50:12:38 | 11.1 ± 1.0 | 7.44 ± 0.40 | 0.151 ± 0.021 |

**Supplementary Table 5.** Permeability coefficients for water, formic acid, and lactic acid in liposomes composed of mixtures of DOPE:DOPC:DOPG or POPE:POPC:POPG in different ratios.

### Supplementary Figures

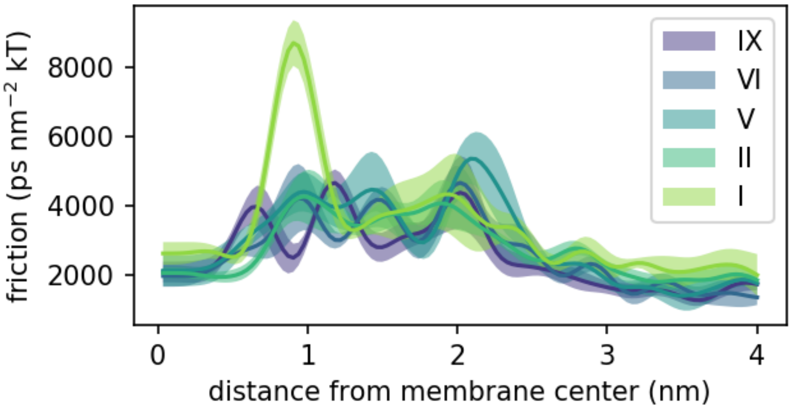


**Supplementary Figure 1.** Friction profiles of solutes of different hydrophobicity levels as a function of the distance from membrane center. More hydrophilic solutes (*i.e.,* level I) exhibit larger friction at the region, where the permeating solute desolvates (around 1 nm).

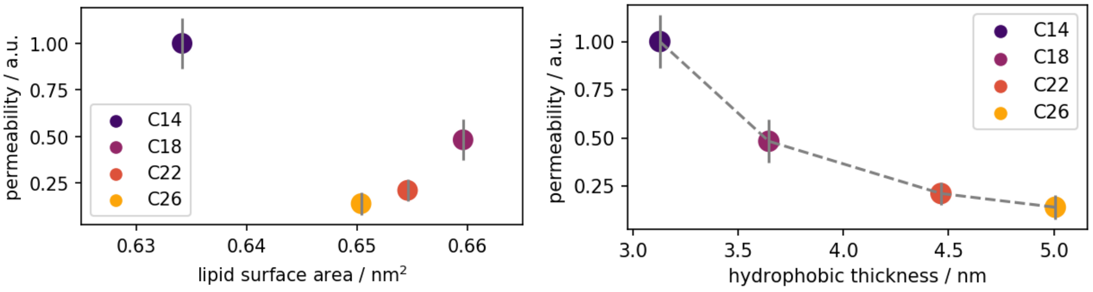


**Supplementary Figure 2.** Permeability coefficient normalized to the membrane with the lowest thickness (C14) as a function of lipid surface area (left) and the membrane hydrophobic thickness (right). While the area per lipid is found to correlate with the permeability coefficient in previous work[1], our simulations do not reveal such correlations. Instead, the expected correlation with thickness is recovered in both experiments and simulations (also in Fig. 2 in the main text).

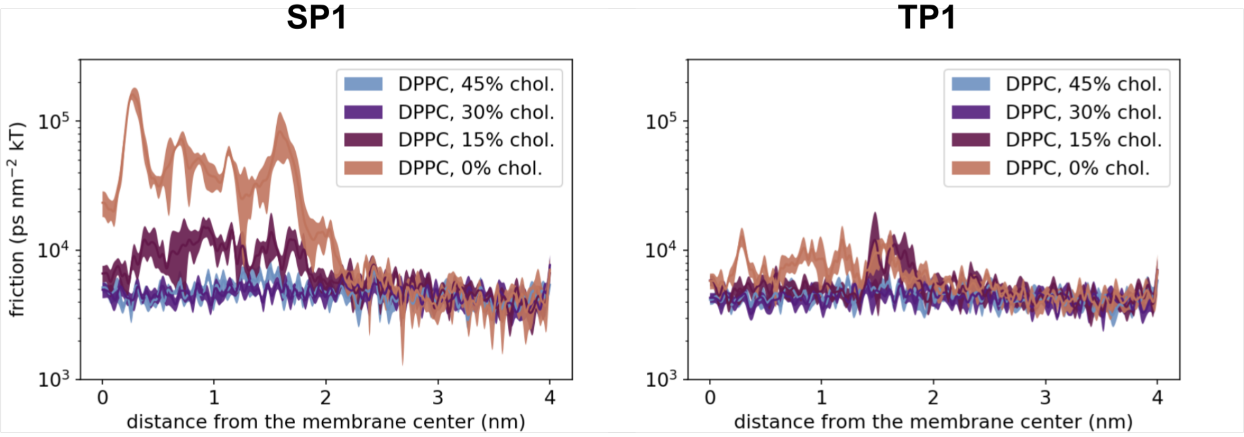


**Supplementary Figure 3.** Friction profiles from MD simulations with particles at the hydrophobicity level III of different sizes – "small" SP1 (size ~ 3 water molecules) and with "tiny" TP1 (size ~ 2 water molecules). The larger solute, SP1, has lower diffusivity through the membranes at a $L_\beta$ phase (DPPC bilayer). Adding 15 mol% of cholesterol to the DPPC membrane leads to a decrease of the friction due to the perturbed packing of the phospholipid tails by the sterols. Such an effect is not as significant for smaller solutes as seen from the changes in the friction profiles from the TP1 solute.

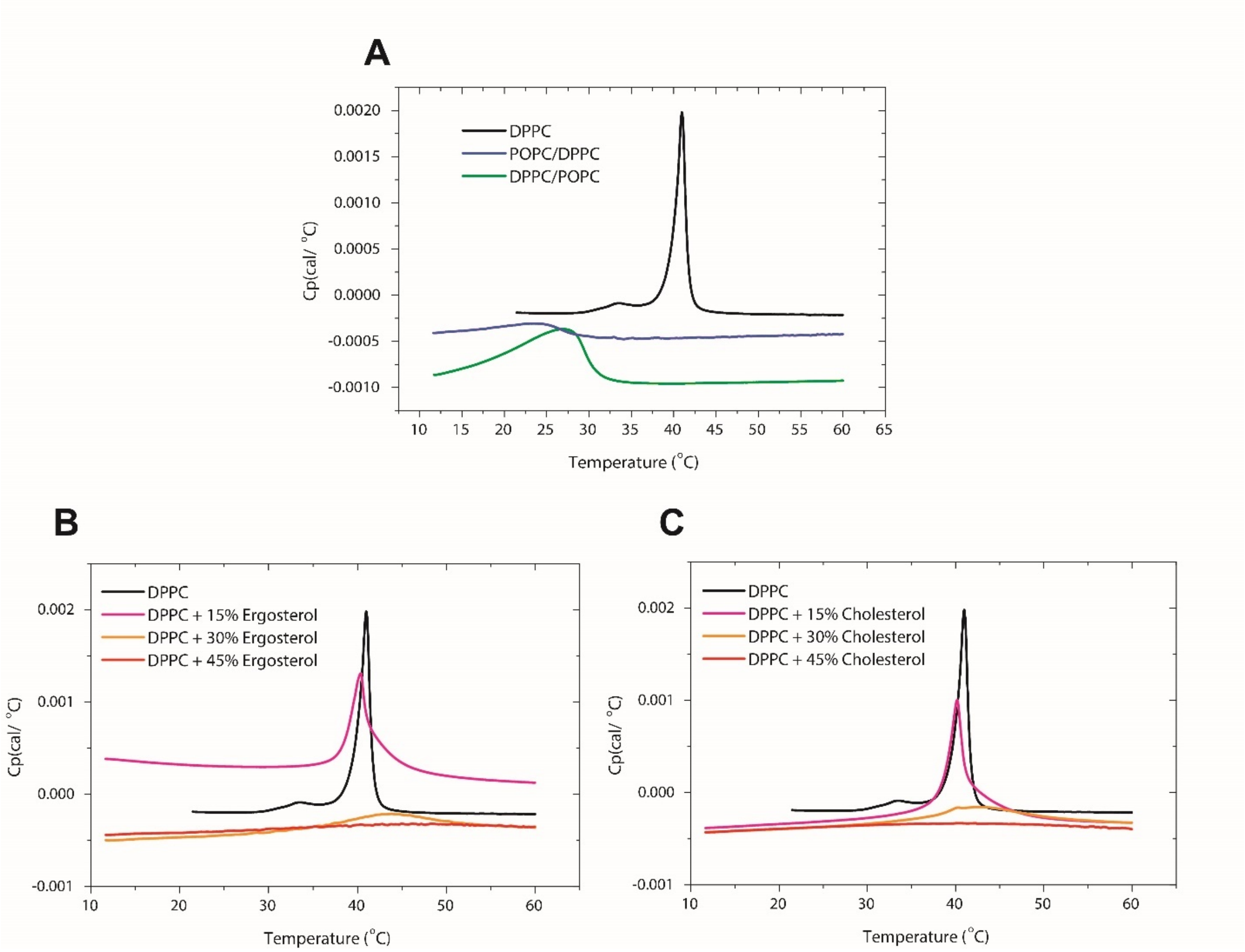


**Supplementary Figure 4.** Normalized DSC endotherms of lipid vesicle solutions, using the baseline as the reference (100 mM KPi buffer pH 7.0). A. Endotherms of pure DPPC (green), POPC/DPPC (ratio 67:33) liposomes (hot pink), and DPPC/POPC (ratio 67:33) vesicles (orange). B. Endotherms of pure DPPC (black) and DPPC + cholesterol vesicles in the ratio of 85:15 (hot pink), 70:30 (green) and 55:45 (cyan). C. Endotherms of pure DPPC (black) and DPPC + ergosterol vesicles in the ratio of 85:15 (hot pink), 70:30 (green) and 55:45 (cyan).

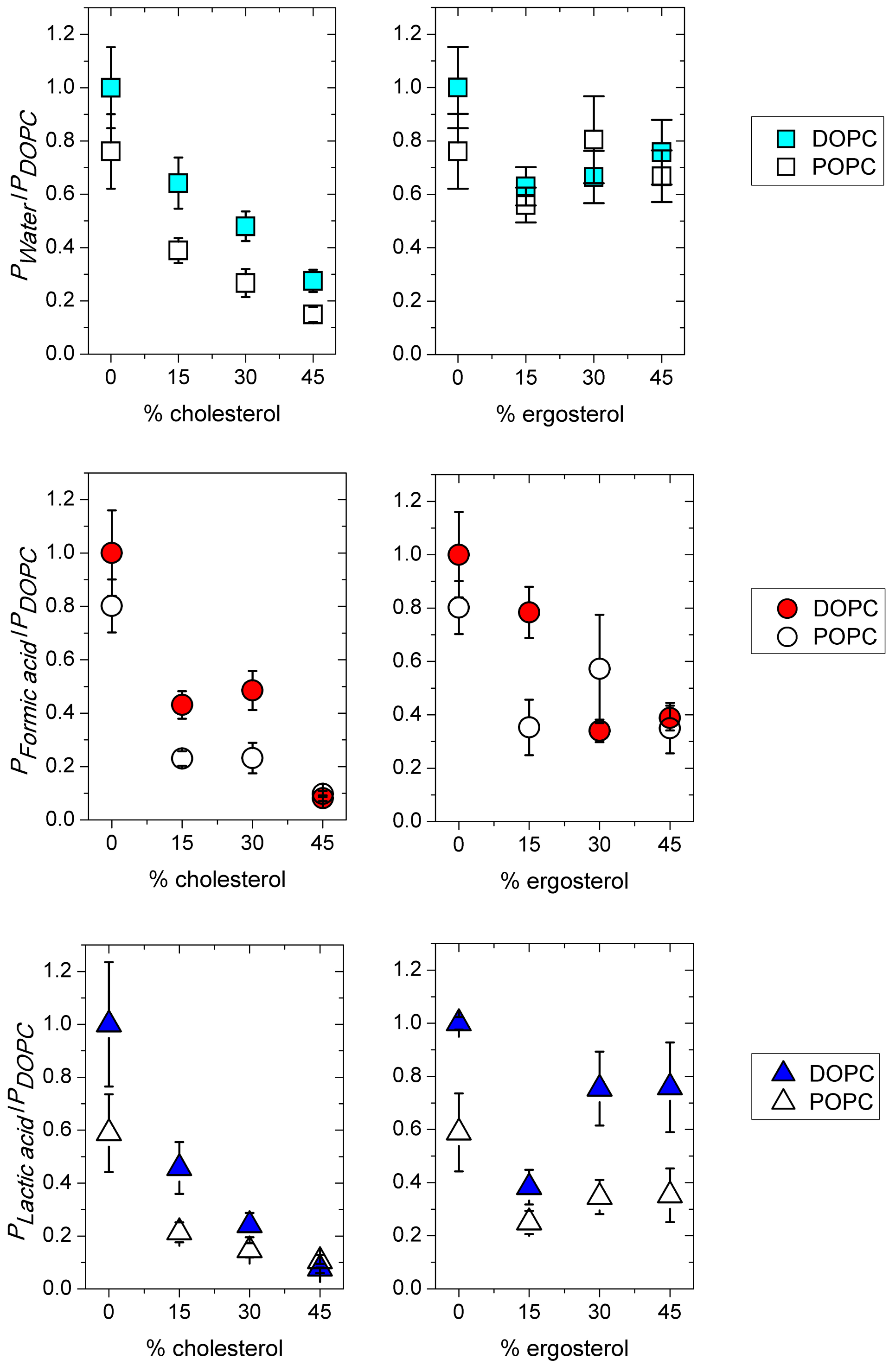


**Supplementary Figure 5.** Permeability of water (squares), formic acid (circles) and lactic acid (triangles) as a function of sterol content in vesicles composed of pure DOPC and POPC, represented by full and hollow markers, respectively. Permeability coefficients *P* (cm/s) are normalized to the value of pure DOPC. The permeability coefficient in DOPC vesicles was 16.0 (±1.7) x $10^{-3}$ cm/s, 6.73 (±0.74) x $10^{-3}$ cm/s, and 0.198 (±0.033) x $10^{-3}$ cm/s for water, formic acid, and L-lactic acid, respectively.

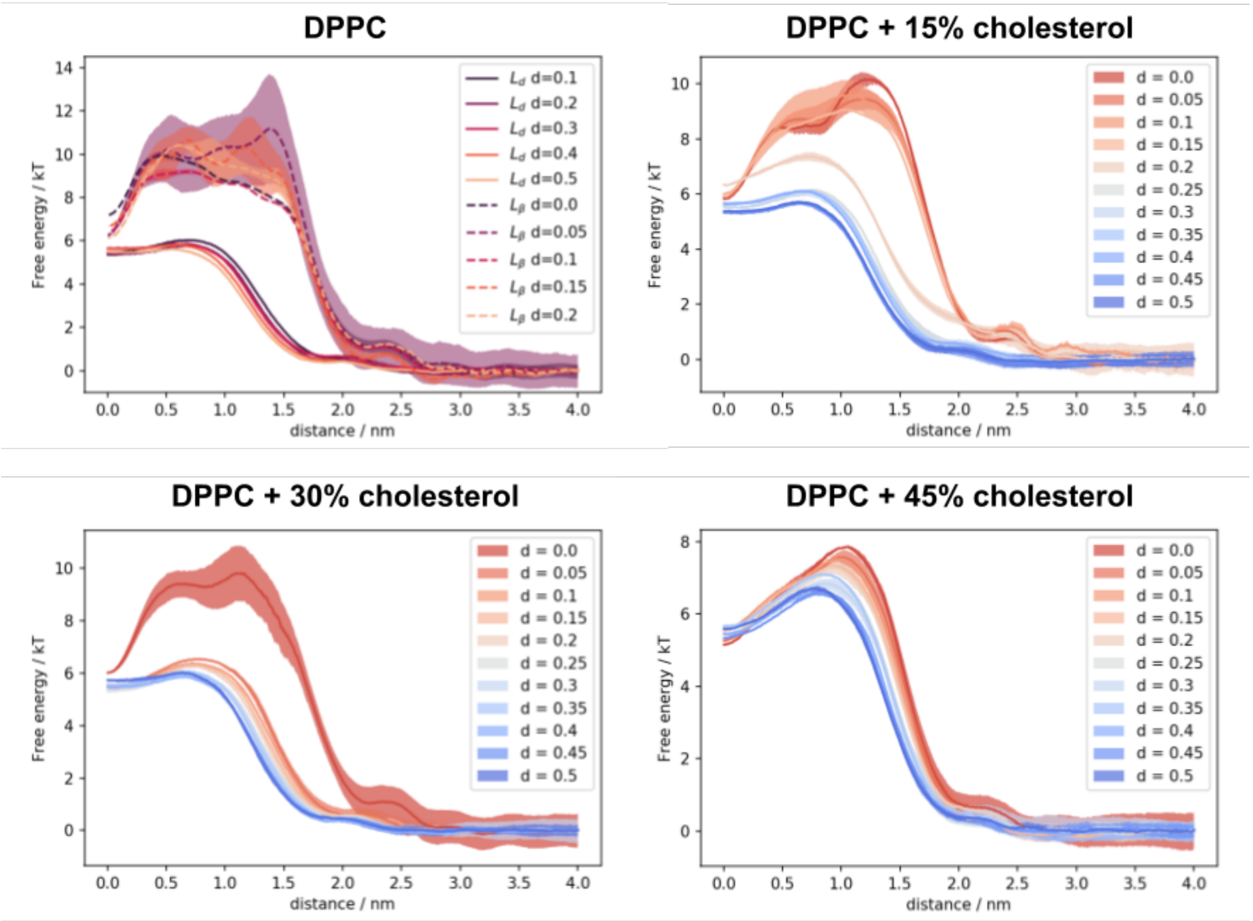


**Supplementary Figure 6.** Free energy profiles from simulated membranes with 0, 15, 30 and 45 mol% cholesterol and a variable unsaturation index *d*. The profiles smoothly change from the composition with POPC (blue, d=0.5) to that with DPPC (red, d=0.0) within a single phase, but exhibit large non-smooth changes with phase transitions affecting the permeability coefficients.

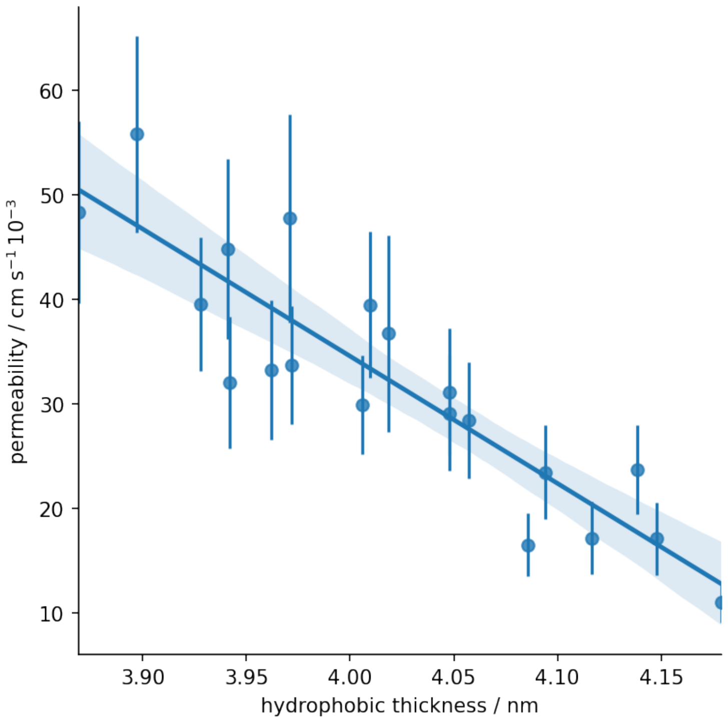


**Supplementary Figure 7.** Regression fit of the permeability as a function of the membrane hydrophobic thickness. Data points are from simulated membranes with a varying degree of tail saturation (index between 0.35 and 0.5) and sterol concentrations (between 0 and 45 mol%). Shaded area denotes the 95% confidence interval.

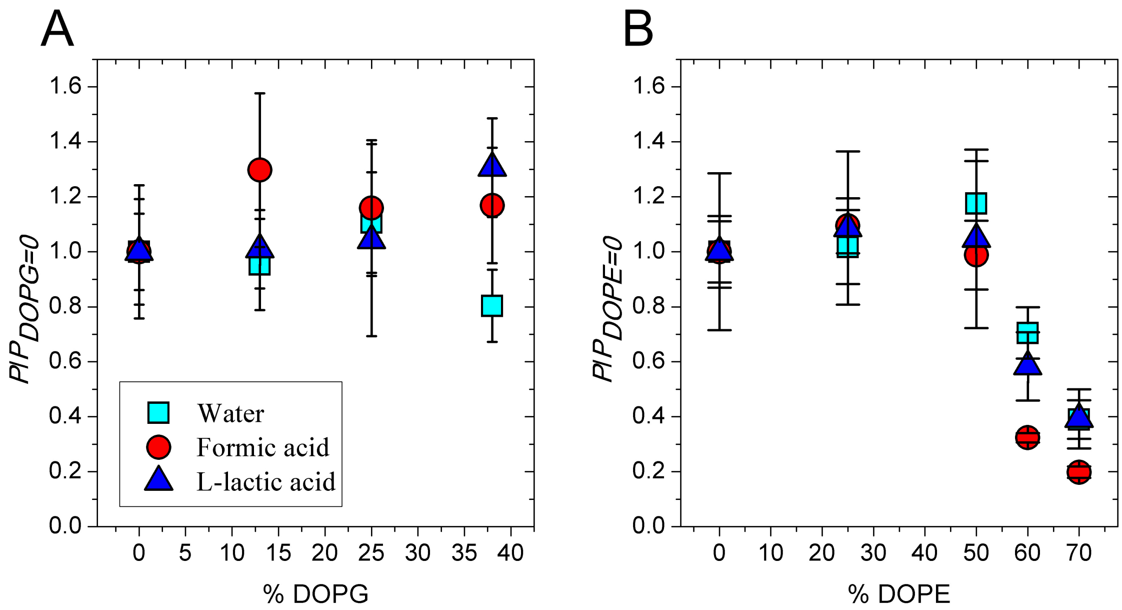


**Supplementary Figure 8.** Permeability of water (cyan squares), formic acid (red circles), and L-lactic acid (blue triangles) as a function of lipid head group composition. A. DOPG titration. The fraction of DOPE was kept constant at 50% and DOPG was varied reciprocally with DOPC. Permeability coefficients are normalized to $P_{DOPG=O}$. The permeability coefficient in $P_{DOPG=O}$ vesicles was 13.8 (±1.9) x $10^{-3}$ cm/s, 6.37 ± (1.09) x $10^{-3}$ cm/s, and 0.116 (±0.016) x $10^{-3}$ cm/s for water, formic acid, and lactic acid, respectively. B. DOPE titration. The fraction of DOPG was kept constant at 25%, and DOPE was varied reciprocally with DOPC. Permeability coefficients are normalized to $P_{DOPE=O}$. The permeability coefficient in $P_{DOPE=O}$ vesicles was 13.0 (±1.2) x $10^{-3}$ cm/s, 7.47 ± (0.83) x $10^{-3}$ cm/s, and 0.119 (±0.024) x $10^{-3}$ cm/s for water, formic acid, and lactic acid, respectively. The numerical values of the permeability coefficients are presented in Supplementary Table S5.

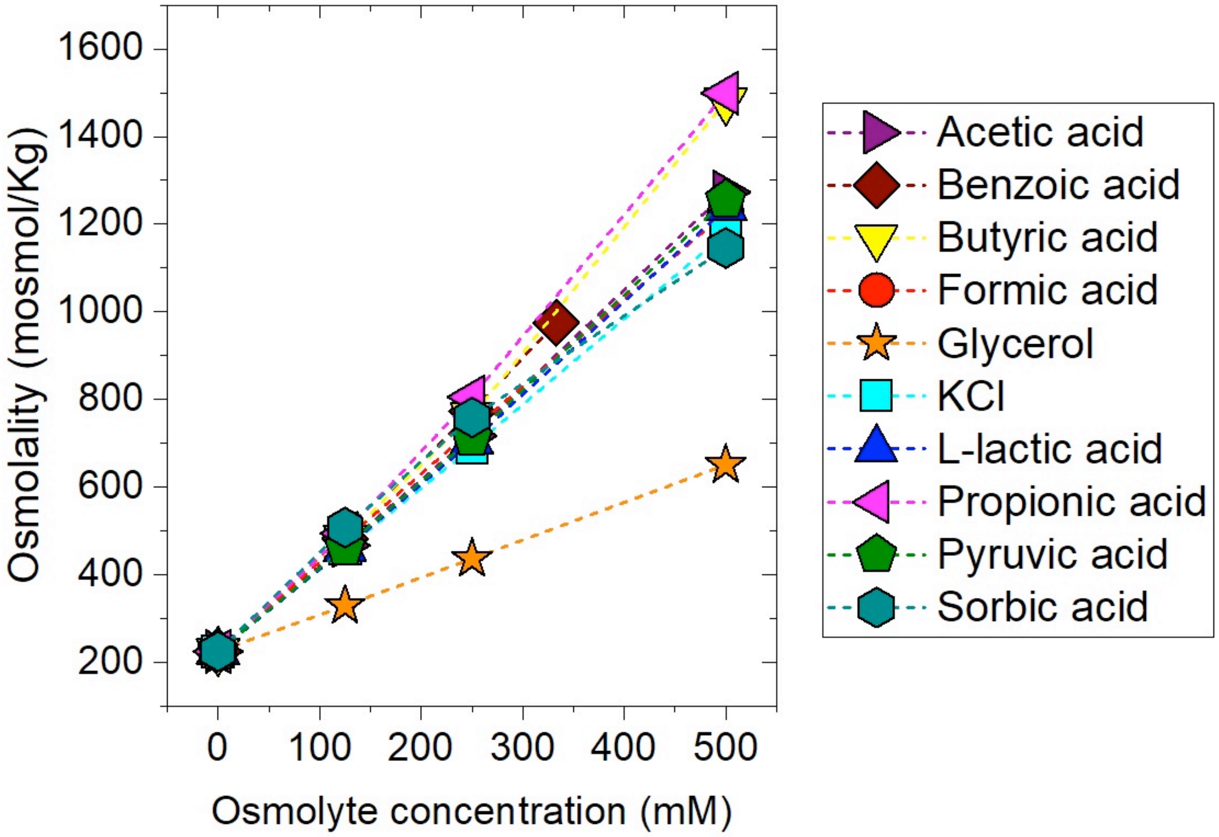


**Supplementary Figure 9.** Plots of the measured osmolality as a function of the osmolyte concentration. The data were fitted with a linear relation (y = mx + q) that was later used to prepare the osmolyte solutions at the desired osmolality of ca. 300 mosmol/kg. The following slopes (m) were used for the calculations: 0.852 (glycerol), 1.822 (K-sorbate), 1.910 (KCl), 2.102 (Na-acetate), 2.367 (Na-benzoate), 2.528 (Na-butyrate), 2.001 (Na-formate), 2.030 (Na-L-lactate), 2.568 (Na-propionate) and 2.068 (Na-pyruvate). The intercept was q = 225.5, which corresponds to the osmolality of the assay buffer (100 mM potassium phosphate, pH 7.0).

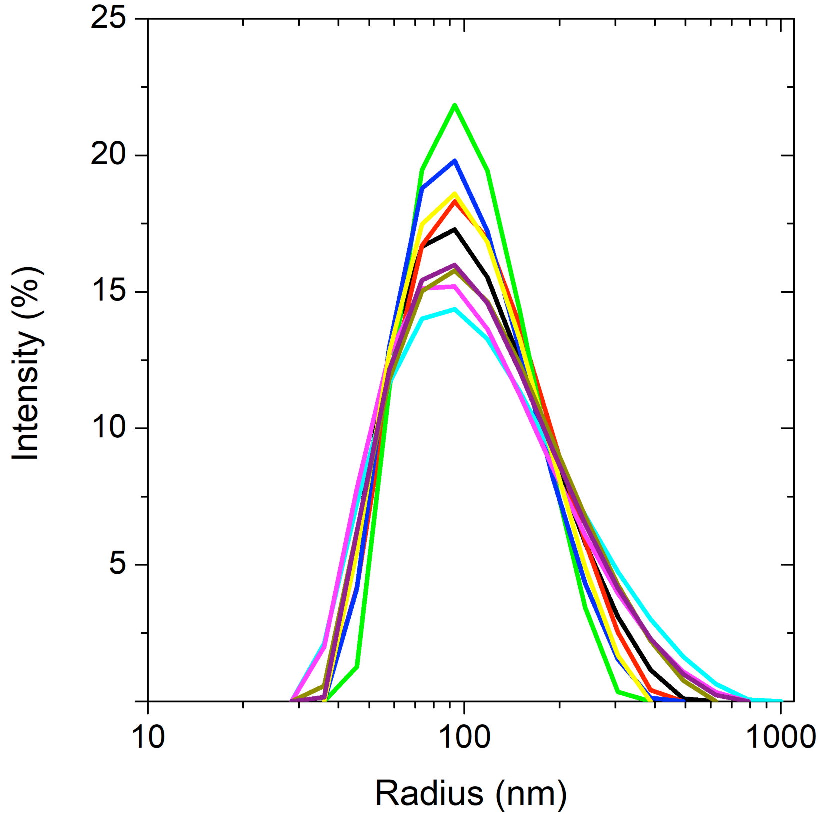


**Supplementary Figure 10**. Size distributions measured with DLS of DOPC vesicles extruded through a 200 nm polycarbonate filter. Ten overlaid acquisitions are shown.
